# Exploration of the single-cell transcriptomic landscape identifies aberrant glomerular cell crosstalk in a murine model of WT1 kidney disease

**DOI:** 10.1101/2022.10.11.511555

**Authors:** Jennifer C Chandler, Daniyal J Jafree, Saif Malik, Gideon Pomeranz, Mary Ball, Maria Kolatsi-Joannou, Alice Piapi, William J Mason, Adrian S Woolf, Paul J Winyard, Andrew S Mason, Aoife M Waters, David A Long

## Abstract

The glomerulus mediates kidney ultrafiltration through specialised epithelial cells called podocytes which line a basement membrane shared with blood capillary endothelium. Cell-cell crosstalk is critical for glomerular function, but its investigation in childhood glomerular diseases has received little attention. *WT1* encodes a transcription factor expressed in podocytes, whose heterozygous variants cause devastating kidney disease in childhood. We used single-cell RNA sequencing and ligand-receptor interaction analysis to resolve the glomerular transcriptional landscape of mice that carry an orthologous human mutation in WT1 (*Wt1*^*R394W/+*^). Podocytes were the most dysregulated cell type in early disease, with disrupted angiogenic signalling preceding glomerular capillary loss. Comparative analyses with additional murine and human glomerular disease datasets identified unique transcriptional changes in WT1 glomerular disease, reflecting a non-immunological pathology, whilst revealing a common injury signature across multiple glomerular diseases. Collectively, this work advocates vascular-based therapies over immunosuppressive drugs in the treatment of WT1 glomerular disease.

## Introduction

Glomeruli are multicellular units in the mammalian kidney that ultrafilter blood, removing toxic components to be excreted in the urine while retaining blood cells and large proteins. This life supporting function relies on a healthy blood supply through glomerular capillaries and the integrity of the glomerular filtration barrier. The filtration barrier is comprised of specialised epithelia called podocytes which wrap around a fenestrated capillary endothelium, sitting on the glomerular basement membrane (GBM). This specialised system is supported by a core of pericyte-like mesangial cells which provide mechanical stability for the glomerular capillary loops. Parietal epithelial cells (PECs) line the Bowman’s capsule, enclosing this structurally and functionally distinct niche from the tubulointerstitium.

Diseases of human glomeruli, also known as ‘glomerulopathies’, can cause nephrotic syndrome, where the circulation is depleted of large proteins, such as albumin, due to their loss through the leaky glomerular filtration barrier into the urine. If blood filtration is more severely impaired, the retention of toxic waste products can be fatal and is referred to as end-stage kidney failure^1^. A multitude of genes have been implicated in glomerulopathies, with podocytes being the cell type most commonly involved, expressing at least 50 of the 80 known genes whose variants result in congenital glomerular disease^2^. These discoveries have had positive clinical impacts, providing genetic information for patients and families. However, fundamental molecular questions remain about how such mutations damage podocytes and how this implicates other glomerular cells through physical and paracrine intercellular signalling, resulting in a multicellular injury response^3,4^.

Early-onset childhood congenital glomerulopathies often present histologically as diffuse mesangial sclerosis (DMS), associated obliteration of glomerular capillaries and mesangial expansion^1,2^. These pathological features result in nephrotic syndrome and reduced glomerular filtration, which rapidly progress to end-stage kidney disease. There are no specific drug treatments for congenital glomerulopathies, with many cases being unresponsive to glucocorticoids or second-line immunosuppressive drugs^1,2^, leaving dialysis or transplantation as the only interventions for these patients. Mutations in the podocyte expressed zinc finger transcription factor *Wilms tumour 1* (*WT1*), account for ~15% of genetically diagnosed congenital glomerular diseases^2^. WT1 governs many key molecular pathways involved in podocyte health and disease, directly regulating approximately half of podocyte-specific genes^5^, at least 18 of which have known variants that result in congenital glomerulopathies^6^. Genomic investigation into pairwise interactions between WT1 and its genetic targets^5,6,7^ has advanced our knowledge of the WT1-regulated transcriptome, but we still lack an integrated understanding of how WT1 initiates multi-cellular glomerular decline and how this can be effectively treated.

One approach to explore glomerular pathophysiology in an unbiased manner is to use single-cell RNA sequencing (scRNA-seq). scRNA-seq yields cell-specific transcriptomes, enabling detection of cell type specific molecular changes, the discovery of rare cell types and the investigation of intercellular communication. This avenue has been used to characterise the multi-cellular transcriptional landscape of glomerular health^8^ and disease^9,10^ at single-cell resolution, with an increasing focus on early disease timepoints, towards the goal of discovering initiators of cellular damage^9,10,11^. However, most studies to-date have focussed on adult pathologies, rather than childhood-onset glomerulopathies. To address this need, we examined a clinically relevant, genetic murine model^12^ of WT1 glomerulopathy (*Wt1*^*R394W/+*^). This model carries a heterozygous *WT1* point mutation, resulting in the substitution of a tryptophan for arginine at codon 394 (*WT1* c.1180C>T; p.R394W). The change is orthologous to a hotspot mutation found in Denys-Drash syndrome^13^ (DDS, OMIM #194080), characterised by early onset steroid-resistant nephrotic syndrome and DMS on histology.

Using the *Wt1*^*R394W/+*^ mouse, we have generated a novel glomerular transcriptomic dataset at single-cell resolution, exploring the early stages of disease. Using cell-specific differential expression and ligand-receptor interaction analyses, we identified early transcriptional characteristics of mutant podocytes, including disrupted angiogenic signalling which precedes glomerular capillary loss. From this, we took a comparative approach with other murine scRNA-seq and human glomerular microarray datasets, identifying transcriptional changes unique to WT1 glomerulopathy, as well as a common injury signature across multiple glomerular pathologies.

## Results and Discussion

### Time course of disease progression in *Wt*^*1R394W/+*^ mice

The glomerulus is a highly specialised multicellular niche (**Fig. 1a**) that becomes structurally and functionally compromised in WT1 glomerulopathy. Prior work has shown that *Wt1*^*R394W/+*^ mouse kidneys appear histologically normal at birth and 3 weeks of age, with proteinuria present by 8 weeks^12^. To further investigate the relationship between renal function and structure we compared histology with urinary albumin/creatinine ratio (ACR) and blood urea nitrogen (BUN) levels; respective indicators of glomerular filtration barrier integrity and renal excretion. At 4 weeks of age, in the absence of histological pathology of DMS (**Fig. 1b**), urinary ACR was significantly elevated (log_10_ ACR 3.96±0.08 µg/mg; *p<0*.*0001*) in *Wt1*^*R394W/+*^ mutant mice compared with *Wt1*^*+/+*^ wild-type littermates (1.91±0.05; **Fig. 1c**), indicating impairment of glomerular filtration barrier functionality. In contrast, BUN levels were similar between *Wt1*^*+/+*^ (45.97±1.65 mg/dL) and *Wt1*^*R394W/+*^ mice (50.74±2.60 mg/dL) at 4 weeks (**Fig. 1d**). By 8 weeks of age, ACR remained significantly elevated in *Wt1*^*R394W/+*^ mice (3.48±0.15 µg/mg;) compared with *Wt1*^*+/+*^ animals (2.21±0.12 µg/mg; *p* < 0.0001, **Fig. 1c**) and histologically, glomeruli showed pathological features of DMS, including global expansion of the mesangial matrix, disorganised vasculature and glomerular tuft fibrosis (**Fig. 1b**). BUN levels were also significantly increased by 8 weeks (*Wt1*^*R394W/+*^, 62.07±2.54 mg/dL *versus Wt1*^*+/+*^ littermates, 48.45±1.81 mg/dL; *p* = 0.0006, **Fig. 1d**). Together this characterisation suggests 4-week-old *Wt1*^*R394W/+*^ mice represent a timepoint when early disease is clinically manifested, detectable through ACR, prior to the emergence of glomerular scaring and deterioration of renal excretory function present by 8 weeks. This rapid time course of disease progression mimics that observed in patients^1,2,13^ and suggests that in *Wt1*^*R394W/+*^ mice, examination of transcriptional changes in the glomerular niche at 4 weeks of age would identify initiators of disease progression.

**Figure 1:**
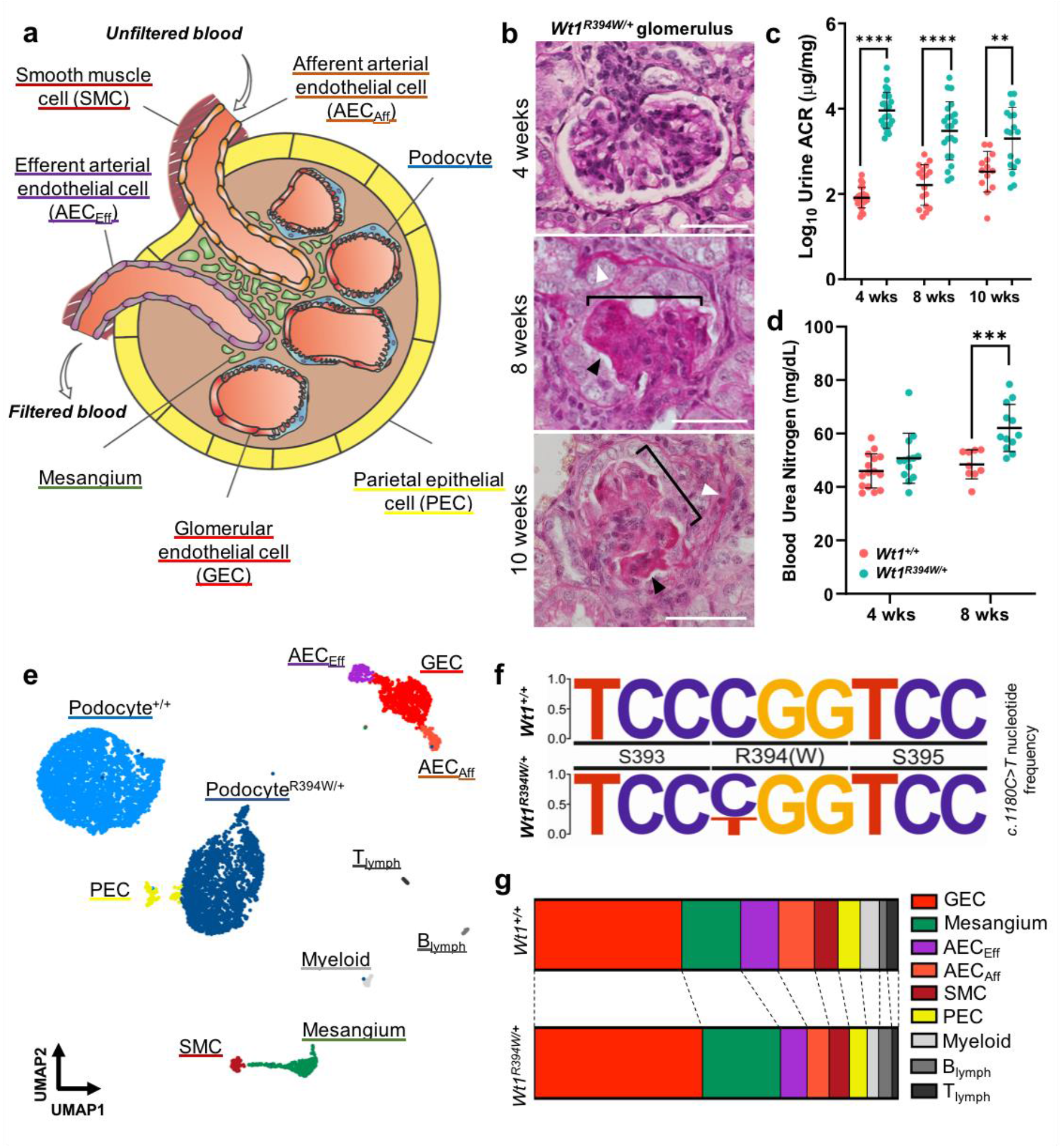
Defining early disease in *Wt1*^*R394W/+*^ mice to generate a glomerular single-cell dataset for WT1 glomerulopathy. a) Schematic of the glomerular niche, comprised of multiple specialised cell types which together, orchestrate plasma filtration, removing toxic waste products for urinary excretion. b) Glomeruli from *Wt1*^*R394W/+*^ mutant kidneys show minimal scarring in early disease at 4 weeks of age, but by 8 weeks, glomeruli show defined features of diffuse mesangial sclerosis, seen as global expansion and scarring of mesangial matrix (black arrowhead), disorganised vasculature (white arrowhead) with shrinking of glomerular tuft (bracketed), these histological features are also seen at 10 weeks; scale bars = 50μm. c) Elevated albumin/creatinine ratio (ACR), indicative of early glomerular damage, is present in *Wt1*^*R394W/+*^ mutant mice (turquoise) from 4 weeks of age, with significantly elevated log_10_ ACR compared to *Wt1*^*+/+*^ littermates (pink) (*t*-test at 4 weeks; *p* < 0.0001; *n=*25, 28; lines represent mean ± SD). d) Blood urea nitrogen (BUN), indicative of renal deterioration, is elevated in *Wt1*^*R394W/+*^ mice by 8 weeks of age (*t*-test; *p* = 0.0006, *n =* 9, 12; lines represent mean ± SD). e) UMAP clustering of sequenced glomerular cells from *Wt1*^*+/+*^ and *Wt1*^*R394W/+*^ mice, a total of 6,846 cell types resolve into eleven transcriptionally distinct cell clusters, of which only podocytes show two distinct clusters based on genotype. f) Plot of *Wt1 c*.*1800 C>T* p. R394W substitution frequency in *Wt1*^*R394W/+*^ mice, showing mutant T allele is at a lower frequency than the wildtype C, at an average of 39% presented. g) Plots showing the proportions of each cell type within each genotype, all non-podocyte cell types are well represented across both *Wt1*^*+/+*^ and *Wt1*^*R394W/+*^.

### Generation of a glomerular single-cell transcriptomic dataset to characterise the onset of WT1 glomerulopathy

To identify changes in glomerular intercellular communication during the initiation of WT1 glomerulopathy, we performed scRNA-seq on glomeruli isolated by Dynabead perfusion at 4 weeks of age in *Wt1*^*+/+*^ and *Wt1*^*R394W/+*^ mice. Using transcardial bead perfusion, glomeruli were isolated (**Supplementary Fig. 1a**) and single-cell suspensions were generated from biochemically representative mice (**Supplementary Fig. 1b-c**), using a protocol^9,14^ further optimised to maximise podocyte viability and yield (see **Methods**). Live cells were isolated using the 10X Genomics Chromium platform and sequenced to a depth of 500 million reads per sample. Putative doublets were removed, as were cells with low feature recovery, or a high proportion of mitochondrial transcripts, generating an aggregated atlas of 6846 cells. Unsupervised clustering and dimension reduction discriminated eleven transcriptionally distinct cell identities (**Fig. 1e**), defined by canonical markers^9,15,16,17^, including two *Wt1*^*+*^ *Nphs2*^*+*^ podocyte clusters (5215 cells), *Emcn*^*+*^ *Ehd3*^*+*^ glomerular capillary endothelium (721 cells), *Ptn*^+^ *Pdgfrb*^*+*^ mesangium (317 cells), *Sox17*^+^ arterial endothelium, including *Plvap*^*+*^ efferent arterioles (141 cells) and *Edn1*^*+*^ afferent arterioles (129 cells), *Acta2*^*+*^ *Myh11*^*+*^ smooth muscle (99 cells) and *Pax8*^+^ *Cldn1*^*+*^ PECs (78 cells). Leukocyte subsets were also identified, including a *Lyz2*^*+*^ myeloid cluster (61 cells), *Cd79a*^+^ *Igkc*^*+*^ B lymphocytes (48 cells) and *Cd3*^*+*^ *Trbc2+* T lymphocytes (37 cells) (**Supplementary Fig. 1d**).

Next, we confirmed assigned genotypes through the alignment of each sample to the *Wt1* primary transcript, demonstrating that the *C>T* mutated allele was expressed as an average of 39% of total *Wt1* coverage in *Wt1*^*R394W/+*^ mice (**Fig. 1f**); in line with the original description of this model^12^. To examine the proportion of cell types from each genotype, we split cells by genotype within each cluster (**Fig. 1g; Supplementary Fig. 1e**), revealing cells from both *Wt1*^*+/+*^ and *Wt1*^*R394W/+*^ mice in all glomerular cell type clusters in similar proportions (**Supplementary Fig. 1f**), with the clear exception of podocytes. Instead, the podocytes from *Wt1*^*+/+*^ and *Wt1*^*R394W/+*^ glomeruli resolved into distinct clusters in two-dimensional space, each corresponding to *Wt1* genotype (**Fig. 1e, Supplementary Fig. 1e-f**). In summary, our scRNA-seq atlas for childhood glomerulopathy contains a similar diversity of cell types compared with adult disease datasets^9,10,11^ and illustrates the extent of podocyte transcriptional disruption driven by a clinically relevant *Wt1* mutation.

### Dysregulation of the podocyte transcriptome dominates early WT1 glomerulopathy

To examine cell type specific changes associated with early WT1 glomerulopathy, we identified differentially expressed genes across the glomerular tuft in podocytes, glomerular endothelial cells, the mesangium and PECs (**Fig. 2a**). Of the 3710 captured genes expressed in podocytes, 268 genes (7.22%) were significantly (log_2_ fold-change (log_2_FC) of ≥ 0.25 and adjusted *p* < 0.05) downregulated and 198 upregulated (5.34%) in *Wt1*^*R394W/+*^ mice compared with *Wt1*^*+/+*^ animals (**Fig. 2a**). Conversely in glomerular endothelial cells, 25 (3.48%) genes were downregulated and 33 (4.5%), from a total of 719 genes detected. Similarly, comparatively low numbers of differentially expressed genes were observed in mesangial cells (12 genes; 1.76% downregulated and 25 genes; 3.66% upregulated) and PECs (6 genes; 1.54% upregulated) (**Fig. 2a, Supplementary Table 1**). To assess whether these transcriptional changes were caused directly by the aberrant binding of mutant WT1, we used CiiiDER^18^ to predict the presence of WT1 binding motifs in all differentially expressed podocyte genes. We found 125 (46.64%) of the podocyte genes downregulated and 107 (54.04%) of those upregulated contained a predicted WT1 regulatory element within 1 kb downstream and 500 base-pairs upstream^6^ of the transcriptional start site. Thus, transcriptional changes in *Wt1*^*R394W/+*^ podocytes are a dominant feature of early WT1 glomerulopathy, partly due to direct transcriptional disruption.

**Figure 2:**
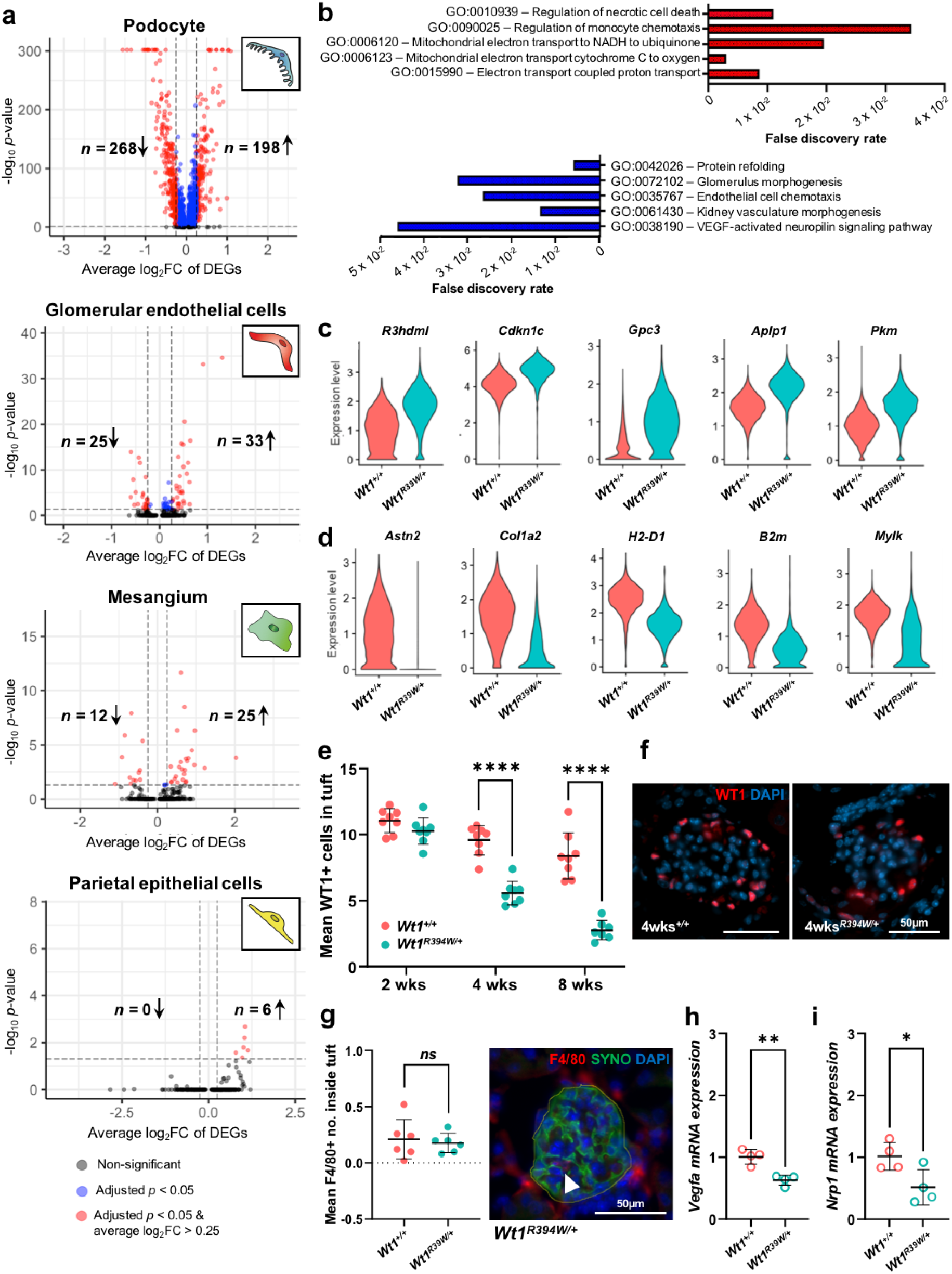
Cell-specific differential expression analyses in the glomerulus shows podocytes as the most affected cell in early WT1 glomerulopathy. a) Volcano plots showing differentially expressed genes in *Wt1*^*R394W/+*^ podocytes (significant hits show average log_2_FC > 0.25 and adjusted *p*-value < 0.05) for podocytes, glomerular endothelial cells, mesangial cells and parietal epithelial cells, showing podocytes as having the highest number and proportion of differentially expressed genes. b) Gene Ontology pathway analyses of podocyte differentially expressed genes, showing the top five upregulated (red) and downregulated (blue) pathways associated with altered gene expression. c) Violin plots of the top five upregulated and D) downregulated genes in *Wt1*^*R394W/+*^ podocytes. e-f) Podocyte (WT1^+^) cell counts in glomeruli (averaged over 50 glomeruli per animal) at 2, 4 and 8 weeks of age show a significant decline in podocyte number from 4 weeks of age (*t*-test; *p <* 0.0001, *n =* 8 mice per group), further reduced by 8 weeks (*t*-test; *p <* 0.0001, *n =* 8 mice per group). g) Myeloid (F4/80^+^) cell counts within the tuft of 4-week-old glomeruli, (averaged over 50 glomeruli per animal) show no difference between *Wt1*^*+/+*^ and *Wt1*^*R394W/+*^ mice (*t*-test; *p =* 0.6864, *n =* 6 mice per group). h) RT-qPCR quantified transcript levels of vascular endothelial growth factor A (*Vegfa*) in *Wt1*^*+/+*^ and *Wt1*^*R394W/+*^ primary podocytes isolated from mice at 4 weeks of age, show a significant decline in *Vegfa* in *Wt1*^*R394W/+*^ podocytes (*t*-test; *p =* 0.0021; *n =* 4 mice per group). i) RT-qPCR quantified transcript levels of Neuropilin-1 (*Nrp1*) in *Wt1*^*+/+*^ and *Wt1*^*R394W/+*^ primary podocytes isolated from mice at 4 weeks of age, show a significant decline in *Nrp1* in *Wt1*^*R394W/+*^ podocytes (*t*-test; *p =* 0.0333; *n =* 4 mice per group).

To identify the functional roles of differentially expressed genes in *Wt1*^*R394W/+*^ podocytes, we conducted Gene Ontology (GO) analysis (**Fig. 2b**). The top upregulated pathways in podocytes suggested metabolic disturbance (GO:0006120 - *mitochondrial electron transport to NADH to ubiquinone*; GO:0006123 - *mitochondrial electron transport cytochrome c to oxygen*; and GO:0015990 - *electron transport couple proton transport*) and cell death (GO:0010939 - *regulation of necrotic cell death*). The latter is in contrast with the proposed function of several of the top five upregulated genes in *Wt1*^*R394W/+*^ podocytes (**Fig. 2c**; **Supplementary Table 1**), which seem to adopt a protective response in glomerular injury. The peptidase inhibitor *R3hdml* (log_2_FC = 1.09) is expressed by podocytes and its overexpression ameliorates TGF-β-induced apoptosis^19^. Fittingly, *R3hdml* knockout in mice show elevated albuminuria, podocyte foot process effacement and GBM thickening. Pyruvate kinase (*Pkm*, log_2_FC = 0.87) overexpression in podocytes protects against mitochondrial dysfunction in murine diabetic nephropathy^20^ and conversely, its loss worsens albuminuria and podocyte injury in an adriamycin model of focal segmental glomerulosclerosis (FSGS)^21^. *Cdkn1c* (log_2_FC = 1.04), encoding p57, is associated with podocyte proliferation in animal models of podocyte injury^22,23^ and patients with proliferative glomerular disease^24,25^, with upregulation of p57 in cultured podocytes being associated with reduced proliferation rates^22^.

Of the downregulated genes in *Wt1*^*R394W/+*^ podocytes, four of the top five associated GO pathways reflected a dampening of angiogenesis (GO:0072102 - *glomerulus morphogenesis;* GO:0035767 - *endothelial cell chemotaxis;* GO: 0061430 - *kidney vasculature morphogenesis*; and GO:0038190 - *VEGF-activated neuropilin signalling pathway*; **Fig. 2b**). Of the top five downregulated transcripts, β2-microglobulin, *B2m* (log_2_FC = −1.07) was classified into the only non-vascular GO term (GO:0042026 - *protein refoldin*g). *B2m* is expressed in cultured mouse podocytes^26^ and is commonly upregulated in patients with glomerulonephritis^27^ (IgA nephropathy and lupus nephritis), with elevated serum levels associated with disease severity in diabetic nephropathy^28^; its downregulation in glomerular disease has not been described. *Col1a2* (log_2_FC = −1.45), encoding collagen α2(I), is present in the glomerular extracellular matrix of healthy mice^29^ and its loss (assessed in *Col1a2*-deficient mice) is associated with sclerotic matrix accumulation in the mesangium, a hallmark of DMS^30^. *Mylk* encodes myosin light chain kinase, involved in stress fibre and focal adhesion formation^31^, processes that are critical for maintaining podocyte architecture, essential for glomerular filtration. Lastly, although the functional role of Astrotactin-2, *Astn2* (log_2_FC = −1.56) in the glomerulus is unknown, a GWAS study identified *ASTN2* as a risk factor locus associated with reduced glomerular filtration rate^32^.

To give a pathological context to these transcriptomic changes, we performed immunofluorescence and RT-qPCR analyses in *Wt1*^*+/+*^ and *Wt1*^*R394W/+*^ mice. To examine the upregulation of pathways of ‘*cell-death*’, we assessed podocyte (WT1^+^) cell number through disease progression, finding glomerular WT1^+^ podocyte cells to be significantly fewer at 4 weeks in *Wt1*^*R394W/+*^ mice (9.58±0.40 cells/glomerular tuft in *Wt1*^*+/+*^ mice *versus* 5.57±0.31 in *Wt1*^*R394W/+*^; *p* < 0.0001, **Fig. 2e-f**), a finding exacerbated by 8 weeks (*Wt1*^*+/+*^, 8.38±0.62 *versus Wt1*^*R394W/+*^, 2.75±0.28; *p* < 0.0001). However, these changes were not detectable in 2-week-old mice (11.04±0.32 and 10.27±0.35 cells/glomerular tuft respectively), reaffirming 4 weeks as an appropriate age to investigate the early stages of glomerular damage. To verify our clustering data showing equal proportions of immune cells in *Wt1*^*+/+*^ and *Wt1*^*R394W/+*^ glomeruli (**Fig. 1g; Supplementary Fig. 1f**) considering the upregulation of GO:0090025 - *regulation of monocyte chemotaxis* in *Wt1*^*R394W/+*^ podocytes (**Fig. 2b**), we examined monocyte influx in 4-week-old *Wt1*^*R394W/+*^ glomeruli, through quantification of cells expressing the monocyte and macrophage marker F4/80^+^ *in situ*. This analysis found similar intraglomerular F4/80^+^ myeloid cell numbers in *Wt1*^*+/+*^ (0.21±0.07 cells) and *Wt1*^*R394W/+*^ mice (0.18±0.04 cells; *p* = 0.6864; **Fig 2g**), which was also the case for extraglomerular F4/80^+^ cell counts (**Supplementary Fig. 1g**). With regard to the pronounced downregulation of angiogenic signalling, RT-qPCR of primary podocytes isolated at 4 weeks of age demonstrated downregulation of both Vascular endothelial growth factor A (*Vegfa*, (*Wt1*^*+/+*^ 1.01±0.06 *versus Wt1*^*R394W/+*^ 0.63±0.04; *p* = 0.0021 **Fig. 2h**) and Neuropilin-1 (*Nrp1, Wt1*^*+/+*^, 1.02±0.11 *versus Wt1*^*R394W/+*^0.52±0.14; *p* = 0.033 **Fig. 2i**) in *Wt1*^*R394W/+*^ podocytes. This accords with previous evidence that WT1 regulates *VEGFA*^6,33^ and given the fundamental role of VEGFA^34^ and NRP1^35,36^ signalling in the maintenance of a healthy filtration barrier, suggests impacts of *Wt1*^*R394W/+*^ podocytes across other cells of the glomerulus.

### Characterisation of the glomerular cell-cell communication landscape identifies defective angiogenic signaling in early WT1 glomerulopathy

We next strove to understand how the transcriptional changes in *Wt1*^*R394W/+*^ podocytes impacts their interaction with other cell types in the kidney glomerulus. To do so we conducted ligand-receptor analysis using the NICHES package^37^. NICHES employs a novel approach that calculates the product of ligand expression on a single cell and the levels of its putative receptor on a receiving cell in a pairwise manner, which, unlike other ligand-receptor analysis tools, generates predicted interactions at single-cell resolution. Interactions were independently screened using STRING^38^ and the literature, such that only validated interactions were included. Firstly, by partitioning our scRNA-seq atlas by genotype, we conducted unsupervised clustering from the NICHES output, to generate interaction maps of the intercellular communication landscape in *Wt1*^*+/+*^ and *Wt1*^*R394W/+*^ glomeruli. Distinct interaction clusters between podocytes and glomerular endothelial cells (**Fig. 3a**), mesangial cells (**Fig. 3b)** and PECs (**Fig. 3c**) were apparent, with no genotype-specific interactions observed. From this, we reasoned that the relative strength of podocyte-derived signals to other glomerular cells could be altered in WT1 glomerulopathy. To assess this, we examined if the most statistically significant interactions, as defined by NICHES, were altered between *Wt1*^*+/+*^ and *Wt1*^*R394W/+*^ glomeruli.

**Figure 3:**
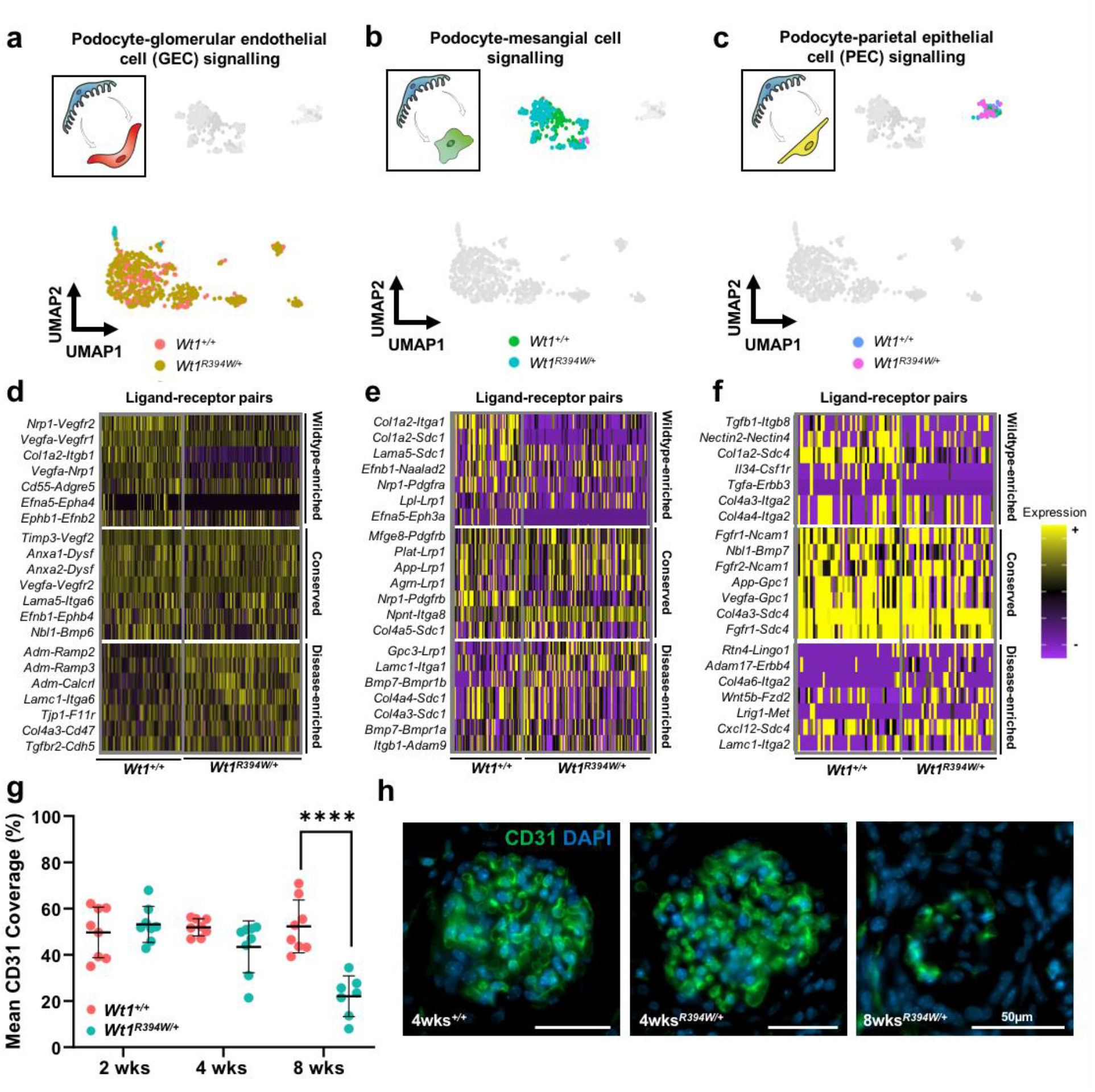
Altered intercellular signalling between *Wt1*^*R394W/+*^ podocytes and endothelium, mesangium and parietal epithelial cells precedes glomerular endothelial loss in WT1 glomerulopathy. Ligand-receptor interaction analysis was performed using the NICHES package^37^ to infer intercellular communication between podocytes and a) glomerular endothelial cells (GEC), b) mesangial cells and c) parietal epithelial cells (PEC). A UMAP enabled visualisation of pairwise interactions between cell types in two-dimensional space, with the following colouring: orange (*Wt1*^*+/+*^ podocytes - *Wt1*^*+/+*^ GEC), yellow (*Wt1*^*R394W/+*^ podocytes - *Wt1*^*R394W/+*^ GEC), green (*Wt1*^*+/+*^ podocytes - *Wt1*^*+/+*^ mesangium), light blue (*Wt1*^*R394W/+*^ podocytes - *Wt1*^*R394W/+*^ mesangium), dark blue (*Wt1*^*+/+*^ podocytes - *Wt1*^*+/+*^ PEC), magenta (*Wt1*^*R394W/+*^ podocytes - *Wt1*^*R394W/+*^ PEC). d-f) Heatmaps of the top seven enriched ligand-receptor interactions by lowest adjusted *p*-value in *Wt1*^*+/+*^ glomeruli, *Wt1*^*R394W/+*^ glomeruli, or shared across both genotypes. g) Endothelial cell (CD31^+^) coverage/glomerular tuft area in glomeruli (averaged over 50 glomeruli per animal) at 2 (*t*-test; *p* = 0.4733; *n =* 8 mice per group), 4 (*t*-test; *p =* 0.0624; *n =* 8 mice per group) and 8 weeks of age, shows a significant decline in CD31^+^ coverage by 8 weeks of age (*t*-test; *p <* 0.0001, *n =* 8 mice per group). h) Representative images of CD31^+^ coverage in the glomerular tuft in *Wt1*^*+/+*^ and *Wt1*^*R394W/+*^ mice at 4 weeks of age with similar endothelial coverage and *Wt1*^*R394W/+*^ at 8 weeks of age when a significant loss in endothelial coverage is observed; scale bars = 50μm.

Between podocytes and glomerular endothelial cells (**Fig. 3d**), *Vegfa-Vegfr2* was amongst the top interactions found in both *Wt1*^*+/+*^ and *Wt1*^*R394W/+*^ mice. The VEGFA signalling pathway, at least partially regulated by WT1, is critical for glomerular function^3,4^, with either overexpression^34,39^ or deletion^34^ of podocyte *Vegfa* resulting in proteinuria, GBM thickening and endothelial cell loss. Annexin-A1 and -A2 to dysferlin signalling (*Anxa1/2-Dysf*), a pro- angiogenic pathway independent of VEGF/VEGFR2^40^ and *Timp3-Vegfr2*, reported to inhibit angiogenesis in human endothelial cells^41^, were also shared between healthy and mutant glomeruli. However, other angiogenic signals involved in the fine-tuning of trans-cellular (between cell) VEGF signalling^42^, including *Nrp1-Vegfr2, Vegfa-Vegfr1* and *Vegfa-Nrp1*, were enriched between podocytes and endothelial cells in healthy glomeruli only (**Fig. 3d**). The loss of these interactions is *Wt1*^*R394W/+*^ glomeruli likely reflects reduction of *Nrp1* and *Vegfa* in mutant podocytes, as neither *Vegfr1* nor *Vegfr2* were differentially expressed in *Wt1*^*R394W/+*^ endothelial cells, with *Nrp1* expression showing a significant increase (log_2_FC = 0.26; **Supplementary Table 1**). Signalling between ephrin ligands and their receptors (*Efna5-Epha4* and *Ephb1-Efnb2*), involved in regulating endothelial cell turnover, migration and permeability^43^, was also present in *Wt1*^*+/+*^ glomeruli, but not enriched in disease. In contrast, the podocyte to endothelial interactions most enriched in *Wt1*^*R394W/+*^ mice involved adrenomedullin and its receptors (*Adm-Ramp2, Adm-Ramp3* and *Adm-Calcr1*), responsible for vasodilatation and vascular barrier function^44^. *Adm* was also detected in the top twenty upregulated (**Supplementary Table 1**) genes in *Wt1*^*R394W/+*^ podocytes and its upregulation has been reported in rodent models of experimental podocyte injury^45,46^.

Intercellular communication analyses also identified known and novel interactions between podocytes and the mesangium. Two interactions (*Mfge8-Pdgfrb* and *Nrp1-Pdgfrb*) conserved across health and disease involved *Pdgfrb*; an established marker of mesangial cells^8,47^ (**Fig. 3e**). *Mfge8* and *Nrp1* regulate the migration of pericytes^48^ and mesenchymal cells^49^ respectively and NRP1-PDGFRβ complex formation is known to regulate the availability of NRP1 to fine-tune VEGFA/VEGFR2 signalling^50^. In contrast, the interaction between *Nrp1* and *Pdgfra* was diminished in *Wt1*^*R394W/+*^ glomeruli. Interactions between podocyte collagen α2(I) and mesangial integrin α1 (*Col1a2-Itga1*) and syndecan-1 (*Col1a2-Scd1)* were also lost in disease, likely driven by the downregulation of *Col1a2* in *Wt1*^*R394W/+*^ podocytes, as neither *Itga1* nor *Scd1* were altered between *Wt1*^*R394W/+*^ and *Wt1*^*+/+*^ mesangial cells (**Supplementary Table 1**). The loss of this interaction highlights a potential molecular mechanism underlying the pathology of *Col1a2* deficiency, associated with DMS in mice^30^. Several podocyte-mesangial interactions enriched in disease involved *Bmp7* signalling, reflecting the significant upregulation of *Bmp7* in *Wt1*^*R394W/+*^ podocytes (**Supplementary Table 1**). As with several of the top upregulated podocyte genes, increased levels of *Bmp7* point towards a protective response, although this time trans-cellular, with elevated BMP7 known to act as a podocyte protective factor in multiple models of mesangial injury^51,52^.

Ligand-receptor analyses also identified interactions between podocytes and PECs, primarily involving secretory pathways. Present in health and disease, these included *Fgfr1/2-Ncam1*, a mediator of FGF signalling^53^ and *Nbl1-Bmp7*, an antagonistic BMP7 interaction, acting on the TGF-β pathway^54^ (**Fig. 3f**). Interactions enriched in *Wt1*^*+/+*^ glomeruli included TGF-β signalling through integrin β8 (*Tgfb1-Itgb8*), required for the release of the mature TGF-β peptide^55^ and *Nectin2-Nectin4*, involved in the trans-cell endocytosis of cytoplasmic cargo^56^, although neither has been investigated in the glomerulus. In *Wt1*^*R394W/+*^ glomeruli, *Adam17-Erbb4* was enriched, which is involved in the regulation of EGF signalling and recently implicated in glomerular injury in murine models of diabetic nephropathy^57^.

As both our GO and ligand-receptor analyses demonstrated a significant disruption to podocyte-derived angiogenic signalling, we analysed glomerular endothelial cell coverage (CD31^+^) through disease progression at 2, 4 and 8 weeks of age. Unlike the onset of podocyte loss, which was present from 4 weeks, endothelial cell number (CD31^+^ coverage/tuft area) was maintained in 2 (*Wt1*^*+/+*^, 49.7±3.7% and *Wt1*^*R394W/+*^, 53.2.±2.8%) and 4 (*Wt1*^*+/+*^, 51.9±1.3% and *Wt1*^*R394W/+*^, 42.9±4.0%) week old glomeruli but showed a ~50% reduction by 8 weeks of age (*Wt1*^*+/+*^, 52.39±4.0% *versus Wt1*^*R394W/+*^, 22.0±3.3%; *p* < 0.0001; **Fig. 3g-h**).

Collectively, our ligand-receptor analysis has highlighted alterations to intercellular communication in the *Wt1*^*R394W/+*^ glomerulus, with a pronounced role for angiogenic signalling that precedes endothelial loss in later disease. By characterising these early changes before the onset of endothelial loss, we have highlighted several podocyte-derived angiogenic molecules, including *Vegfa, Nrp1* and ephrin ligands, associated with subsequent endothelial decline. Two additional *WT1* mutations in human (*WT1*_c.1316G>A; p.R439H)^58^ and mouse (*Wt1*^*R362X/+*^)^59^ have been described with glomerular endothelial damage as a feature of their pathology. Together this suggests vascular focussed therapies may offer a targeted treatment approach for WT1 glomerulopathies.

### Common signatures of podocyte injury across murine and human glomerulopathies

To improve our molecular understanding of WT1 glomerulopathy relative to other glomerular diseases, we conducted an integrated comparison of *Wt1*^*R394W/+*^ glomeruli with three other murine scRNA-seq datasets^9^: 1) the immune-mediated nephrotoxic nephritis model (**Supplementary Fig. 2a**), analogous to adult autoimmune glomerulonephritis; 2) a leptin-deficiency (BTBR *Lep*^*ob/ob*^) model of adult diabetic nephropathy (**Supplementary Fig. 2b**); and 3) a CD2AP-null (*Cd2ap*^*−/−*^) model of congenital FSGS, which also results in compromised immune function due to the role of CD2AP in T-cell adhesion^60^ (**Supplementary Fig. 2c**). Akin to our *Wt1*^*R394W/+*^ dataset, scRNA-seq data from each of these models was taken at timepoints corresponding to the onset of proteinuria, but prior to pathological changes in the glomerulus^9^. Differential expression analyses (log_2_FC of ≥ 0.25 and adjusted *p* < 0.05) were run on *Nphs1*^*+*^ and *Nphs2*^*+*^ podocytes from each model, generating gene lists for comparison across all four disease models (**Fig. 4a-b**). This approach identified a common signature of 23 upregulated and 5 downregulated genes by podocytes across all four models (**Fig. 4c**); with 47 upregulated and 17 downregulated in three or more of the four datasets (**Supplementary Table 2**).

**Figure 4:**
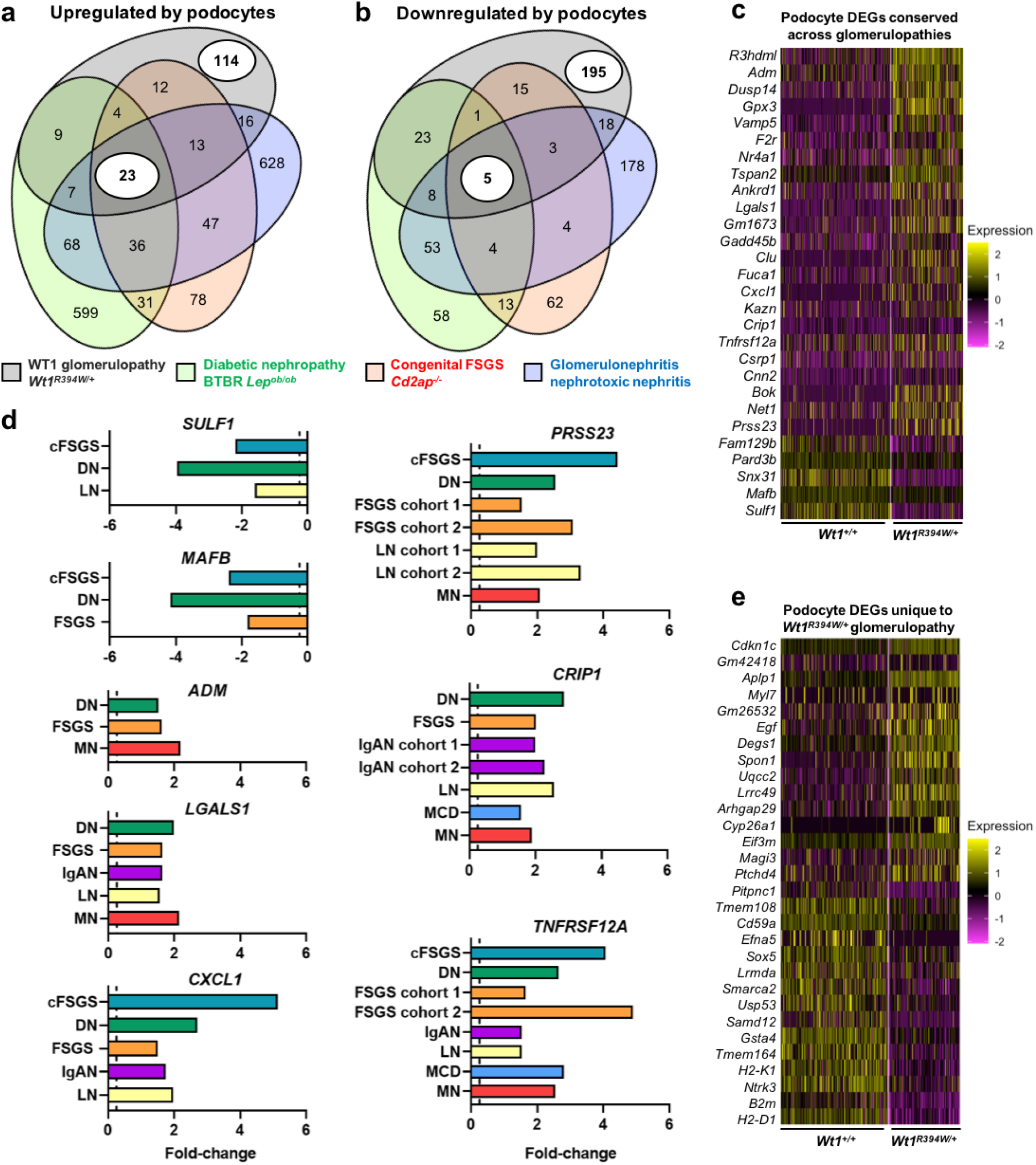
Damaged podocytes share a common signature in both murine and human glomerulopathies in children and adults, but *Wt1*^*R394W/+*^ glomerulopathy has a specific transcriptional profile. a) Venn diagram of upregulated differentially expressed genes (DEGs) in diseased podocytes across four murine models of early glomerular disease (*Wt1*^*R394W/+*^, black; nephrotoxic nephritis, blue; *Lep*^*ob/ob*^ diabetes, green; and *Cd2ap*^*−/−*^, red), highlighting that the majority of *Wt1*^*R394W/+*^ differentially expressed genes are unique, but 23 genes are common across all four glomerulopathies. b) Venn diagram of downregulated podocyte differentially expressed genes across four murine models of early glomerular disease highlighting that the majority of *Wt1*^*R394W/+*^ DEGs are unique, but 5 genes are common across all four glomerulopathies. c) Heatmap of the 23 and 5 conserved podocyte genes in the *Wt1*^*R394W/+*^ dataset that are up or downregulated across four early disease murine glomerular models. d) Bar graphs of the fold change increase or decrease of conserved genes in murine models similarly dysregulated in three or more human glomerular pathologies (microarray data, Nephroseq); abbreviations as follows: HD, healthy donor; cFSGS, collapsing focal segmental glomerulosclerosis glomeruli; DN, diabetic nephropathy glomeruli; FSGS, focal segmental glomerulosclerosis glomeruli; IgAN, IgA nephropathy glomeruli; LN, lupus nephritis glomeruli; MCD, minimal change disease glomeruli; MN, membranous nephropathy glomeruli. e) Heatmap of podocyte DEGs that are unique to *Wt1*^*R394W/+*^ glomerulopathy and were not present in nephrotoxic nephritis, *Lep*^*ob/ob*^ diabetes, or *Cd2ap*^*−/−*^ models.

To explore the relevance of these conserved murine genes to human disease, we used Nephroseq to analyse microarray datasets derived from a spectrum of human glomerular diseases. These included: variants of FSGS^61,62^, classified by focal scarring of the glomeruli; minimal change disease (MCD)^61,62^, showing normal light microscopy; three immune-mediated glomerulopathies, IgA nephropathy (IgAN)^61,63^, membranous nephropathy (MN)^61^ and lupus nephritis (LN)^61,64,65^; and diabetic nephropathy (DN)^61,66^, characterised by glomerular hypertrophy and sclerosis. Thirteen of the 28 (46.43%) transcripts differentially expressed across murine models were also altered in at least one human glomerular dataset (**Table 1**) and eight genes were altered in three or more human glomerulopathies. *SULF1* (**Supplementary Fig. 3ai-iii**) and *MAFB* (**Supplementary Fig. 3bi-iii**) were downregulated and *ADM* (**Supplementary Fig. 3ci-iii**) upregulated in three human disease datasets, *LGALS1* (**Supplementary Fig. 3di-v**), *CXCL1* (**Supplementary Fig. 3ei-v**) and *PRSS23* (**Supplementary Fig. 3fi-vii**) transcripts were significantly elevated in five, *CRIP1* (**Supplementary Fig. 3gi-vi**) upregulated in six and *TNFRSF12A* (**Supplementary Fig 3hi-viii**) increased in seven glomerular disease datasets (**Fig. 4d**).

**Table 1.**
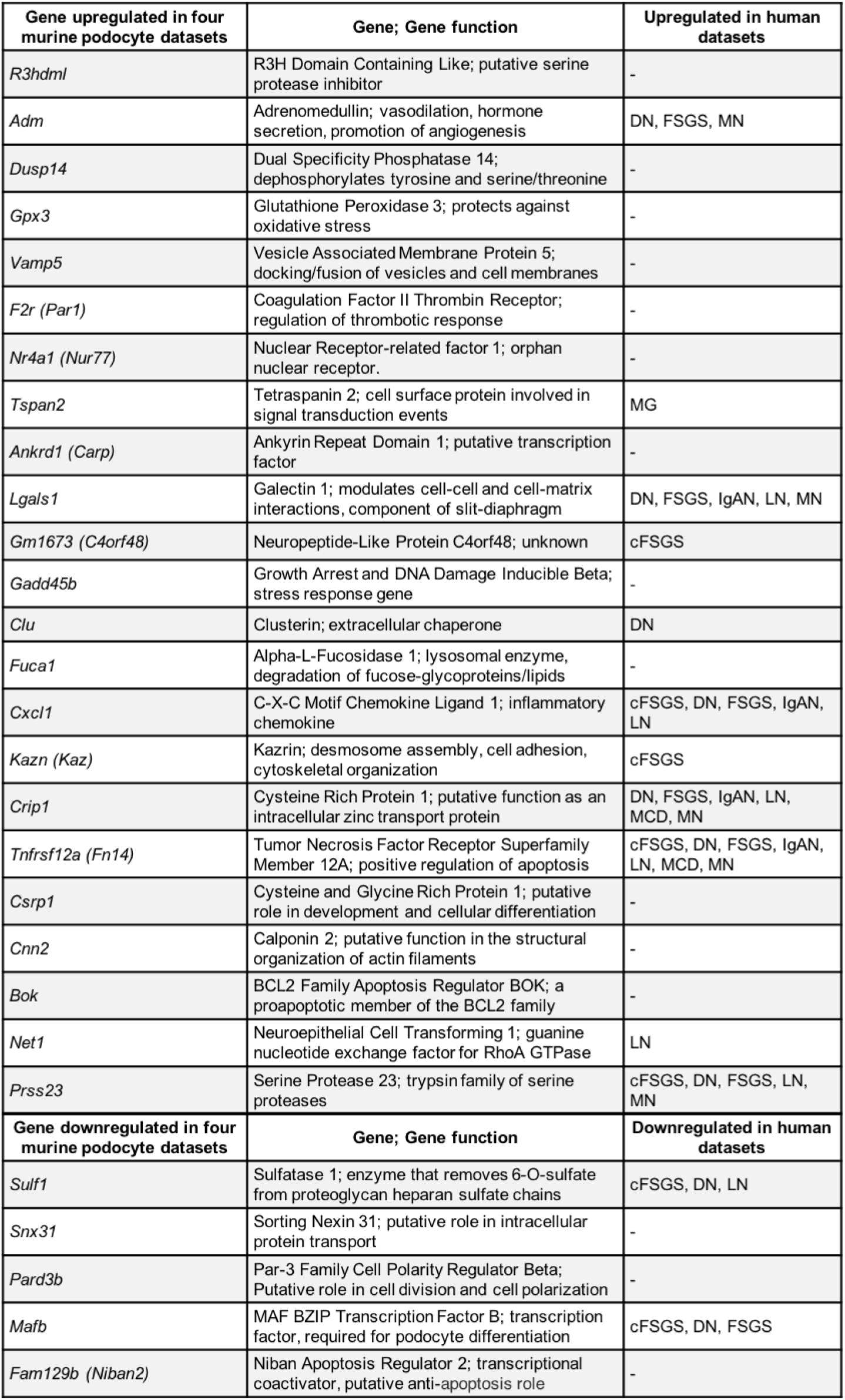
Podocyte genes dysregulated in *Wt1*^*R394W/+*^, nephrotoxic nephritis, *Lep*^*ob/ob*^ diabetes and *Cd2ap*^*−/−*^ murine scRNA-seq datasets, with associated changes in human glomerulopathies. All 23 upregulated and 5 downregulated podocyte genes found in murine early disease datasets^9^ (*Wt1*^*R394W/+*^, nephrotoxic nephritis, *Lep*^*ob/ob*^ diabetes and *Cd2ap*^*−/−*^) a nd their corresponding occurrence in human glomerular pathologies (Nephroseq). Abbreviations as follows: HD, healthy donor; cFSGS, collapsing focal segmental glomerulosclerosis glomeruli; DN, diabetic nephropathy glomeruli; FSGS, focal segmental glomerulosclerosis glomeruli; IgAN, IgA nephropathy glomeruli; LN, lupus nephritis glomeruli; MCD, minimal change disease glomeruli; MN, membranous nephropathy glomeruli.

These seven genes span a variety of functions, *ADM* and *SULF1* are implicated in secretory signalling pathways. *ADM*, enriched in *Wt1*^*R394W/+*^ podocyte to endothelial signalling interactions (**Fig. 3d**), is a secreted vasodilatory peptide. *SULF1*, sulfatase-1, the eighth most downregulated gene in *Wt1*^*R394W/+*^ podocytes (**Supplementary Table 1**), is a modulator of growth factor signalling in the GBM and a direct target of WT1^67^. Interestingly, global *Sulf1*^*−/*^; *Sulf2*^*−/−*^ knock-out mouse, displays proteinuria, glomerulosclerosis and a reduced distribution of VEGFA in the GBM^69,70^, suggesting a role for *SULF1* beyond that of WT1 glomerulopathy alone.

Both *LGALS1* (Galectin-1) and *CRIP1* have more structural roles. Galectin-1 is a podocyte-produced component of the slit-diaphragm and its increased glomerular expression is a feature of childhood DMS and FSGS^71^, as well as adult DN^72^. The expression of *CRIP1* has not been described in podocytes, however it has been defined as a marker of renin cells in the juxtaglomerular region^73^. The enzymatic function of serine protease 23 (*PRSS23*) has also received little attention in the context of glomerular disease, but it is upregulated in human FSGS glomeruli^74^ and in podocytes and mesangial cells from a *Cd2ap*^*+/−*^; *Fyn*^*−/−*^ mouse model of FSGS^75^. The latter study highlighted that PRSS23 can activate Par2 (Protease-Activated Receptor 2), a receptor implicated in TGFβ1-induced podocyte injury. *MAFB*, encoding a transcription factor required for maintenance of podocyte differentiation, is decreased in podocytes from FSGS patients and its overexpression alleviates adriamycin-induced FSGS in mice^75^.

The final two molecules *CXCL1* and *TNFRSF12A*, upregulated in five and seven human glomerular diseases respectively, are both immune factors. CXCL1 is a chemokine commonly implicated in glomerulonephritis, regulating infiltration of neutrophils into the glomerulus^76^. Similarly, activation of the Tumor necrosis factor-like weak inducer of apoptosis (TWEAK) receptor (*TNFRSF12A*), has been shown to promote inflammation and fibrosis in LN^77^ and crescent formation in IgAN^78^. Our findings not only show the upregulation of *CXCL1* and *TNFRSF12A* in glomerulonephritis pathologies, but also across other nephrotic diseases from a non-immune origin. Sanchez-Nino *et al*. have also shown elevated *TNFRSF12A* in human FSGS and MN glomeruli and a non-immune experimental mouse model, associated with elevated *Ccl2*, a potent inflammatory chemokine driving macrophage infiltration^79^. In contrast, despite *CXCL1* and *TNFRSF12A* upregulation in *Wt1*^*R394W/+*^ podocytes, we did not see an upregulation of *Ccl2* (**Supplementary Table 1**), nor an increase in intraglomerular macrophage number (**Fig. 2g**). Together this suggests differential roles for *CXCL1* and *TNFRSF12A* across a broader range of glomerulopathies than currently characterised.

### WT1 glomerulopathy has a unique transcriptional signature

In performing our comparative analyses across murine podocyte scRNA-seq datasets, we have also revealed specific alterations unique to each pathology. These disease-specific transcriptional changes account for the vast majority of differentially expressed genes in each pathology (**Fig. 4a-b**). Focussing on WT1 glomerulopathy, the 114 upregulated (57.3% of the total upregulated) and 195 downregulated (72.8% of the total downregulated) genes unique to *Wt1*^*R394W/+*^ disease point towards its distinct aetiology (**Supplementary Table 2**); the top thirty unique genes (by log_2_FC) are shown in **Fig. 4e**.

With this in mind, we strove to explore a key aetiological hallmark of WT1 glomerulopathy and other congenital glomerular diseases, their resistance to glucocorticoids and second-line immunosuppressive therapies^1,2^. To do so, we interrogated the list of unique differentially expressed genes from each of the four murine podocyte datasets, identifying those classified in the broadest immunological GO term (GO:0002376 - *immune system process*). The unique podocyte gene lists from nephrotoxic nephritis, DN and *Cd2ap*^*−/−*^ models all showed an overall upregulation of genes involved in ‘*immune system processes’*, whilst *Wt1*^*R394W/+*^ was the only model in which a downregulation of this GO term dominated (**Supplementary Table 3**). This bias for the suppression of *immune system processes* in *Wt1*^*R394W/+*^ podocytes is unlikely to reflect direct WT1 mis-regulation, as only 2 of the 14 immune-related genes were predicted to contain a WT1 regulatory element. Together this highlights a critical feature of WT1 aetiology, where a decrease in immune-related genes defines early pathology. This provides a potential explanation for the suboptimal efficacy of immunosuppressive drugs in treating WT1 glomerulopathy in a clinical setting.

## Conclusions

Our results constitute a first scRNA-seq glomerular dataset of a murine model carrying a clinically relevant point mutation observed in patients. Here, we have shown podocytes to be the most altered glomerular cell type in early *Wt1*^*R394W/+*^ pathology, consistent with the localisation of WT1 expression in the postnatal kidney, impacting podocyte cell viability and glomerular paracrine signalling. Crucially, this highlights a window for therapeutic intervention, prior to an irreversible decline in organ function and before other cells in the glomerulus, such as the glomerular endothelium, are damaged through aberrant *Wt1*^*R394W/+*^ podocyte signalling. Furthermore, by integrating our findings with publicly available datasets, we have advanced our understanding of glomerular pathologies across murine models and into human, identifying a signature of molecules that define podocyte injury, which hold promise for novel treatment approaches for multiple glomerular diseases in the future. For WT1 glomerulopathy, this work reaffirms the unfounded use of immunosuppressive strategies in disease management. Instead, it points towards the use of alternatives, such as vascular based therapies to modulate disrupted podocyte cell-signalling and associated loss of the glomerular endothelium that occurs concurrently with decline in kidney function.

## Methods

### Experimental Animals and Procedures

Male C57BL/6 mice heterozygous for *Wt1 c*.*1800 C>T* p. R394W were backcrossed with female MF1s (Charles River) for two generations^12^; second-generation male offspring (*Wt1*^*R394W/+*^) were used for all subsequent analysis. All procedures were approved by the UK Home Office. Renal function was assessed at 4, 8 and 10 weeks of age by urinary albumin/creatinine ratio (ACR) via enzyme-linked immunosorbent assay of albumin (Bethyl Laboratories, Montgomery, TX) and creatinine (Cayman Chemical, 500701). Blood urea nitrogen (BUN) levels were quantified in plasma taken at 4 and 8 weeks of age using the QuantiChrom™ Urea Assay Kit (DIUR-100, BioAssay Systems). Histological analyses were conducted on 7μm periodic acid–Schiff stained sections.

### Glomerular isolation and preparation of single cell suspensions

Prior to glomerular isolation urinary ACR and plasma BUN were analysed across littermates at 4 weeks as described. Representative individuals (*n* = 2) for each of *Wt1*^*+/+*^ and *Wt1*^*R394W/+*^ were used for single-cell RNA-seq. Glomeruli were isolated as previously described^9,14^ with some adaptations to maximise podocyte yield. Due to the smaller size of the mouse at 4 weeks old, limiting the efficacy of inter-arterial bead perfusion, transcardial perfusion was used to increase glomerular yield, mice were perfused with 40 ml of ice-cold Hanks’ Balanced Salt Solution without calcium and magnesium (HBSS^−/−^), containing 300 μL M-450 Epoxy Dynabeads (14011, Invitrogen). Both kidneys were removed and minced into small pieces with a scalpel and incubated in 3 ml prewarmed digestion buffer (1.5 U/ml Liberase TM, 100 U/ml DNase I in HBSS^−/−^) at 37°C for 20 minutes with constant agitation (300 RPM) in a ThermoMixer (Eppendorf). All subsequent steps were performed as previously described^9,14^, with the addition of bovine serum albumin (BSA)-coated plastics throughout to prevent cell adhesion (10% BSA in DPBS^−/−^ and rinsed with DPBS^−/−^).To enhance podocyte number in the final cell fraction, the mechanical digest approach was optimised as follows: gentle pipetting every 10 mins for 30 mins, at 30 minutes each sample was transferred into a 2 ml syringe (without passing via the needle) and slowly and steadily pushed through 25 ½ gauge needle followed by a 27 ½ gauge needle (smaller gauge needles were not used as these were associated with podocyte loss). Samples were then incubated for an additional 5 minutes with one pipette mix. At 45 minutes the digest was stopped with 10 ml ice-cold DBPS^−/−^ with 10% foetal bovine serum (FBS) and placed on a magnetic separator to remove magnetic beads. The supernatant was sieved (100 µm cell strainer) and rinsed with DPBS^−/−^ + 10% FBS to remove aggregates. Cells were resuspended in DPBS^−/−^ + 2% FBS, 5 mM EDTA and underwent a live/dead cell sort with propidium iodide viability dye with 98.1-98.9% viability across all four samples.

### Library preparation, single-cell sequencing and alignment

Healthy and disease glomerular single-cell suspensions were processed in parallel using the Chromium Next GEM Single Cell 3ʹ Reagent Kits v3.1 (Dual Index) kit according to manufacturer’s instructions. GEMs underwent 3’ GEX library preparation and libraries were pooled. All four samples underwent 150 base pair (bp) paired-end read sequencing on one lane of an S4 flow cell (Illumina NovaSeq 6000) to a minimum depth of 500 million reads per sample (Source BioScience, Nottingham, UK). Reads were mapped individually from each sample to the 10x-approved gex-mm10-2020-A mouse reference sequence using 10X Genomics Cell Ranger v5.0.1^80^. Data were separately aligned to the *Wt1* primary transcript using HISAT2 v2.2.1^81^ to confirm the presence of the Wt1p.R394W heterozygous mutation in the two *Wt1*^*R394W/+*^ samples.

### Quality control and pre-processing of single-cell sequencing data

Single-cell sequencing data were aggregated and processed using Seurat (v4.1.1)^82^ in R Studio (v 2022.02.0). The full script for analysis had been made available at https://github.com/daniyal-jafree1995/collaborations/blob/main/Chandleretal_2022_WT1glomerulopathyscRNAseq.

Quality control criteria were empirically determined, including the removal of cells with fewer than 150 RNA features or mitochondrial feature representation greater than 20% and putative doublets (DoubletFinder 2.0.3^83^). 93.77% of cells (6846/7301) were retained in the final count matrix, with levels consistent across each donor. The resulting count matrix was corrected for ambient RNA contamination using the SoupX (v1.6.1) package^84^. Samples were then normalised, scaling expression by all genes detected, before dimension reduction with principal component analysis, which identified 16 informative principal components. This reduced dataset was used for nearest-neighbour graph construction and unsupervised clustering (Louvain method), with a 0.4 resolution selected based on apparent cluster separation on indicative marker genes and visualised by UMAP.

### Cell type identification

Cell type clusters were identified using canonical markers of glomerular cells and informed by published scRNA-seq studies of murine glomeruli, vasculature and immune cells^9,15,16,17^. These markers were visualised using the DotPlot function in Seurat and included *Wt1* and *Nphs2* for podocytes, *Emcn* and *Ehd3* for glomerular endothelial cells, *Ptn* and *Pdgfrb* for mesangial cells, *Sox17* for arterial subsets including *Plvap*-expressing efferent arteriole and *Edn1*-expressed afferent arteriole, *Acta2* and *Myh11* for vascular smooth muscle cells, *Pax8* and *Cldn1* for parietal epithelial cells, *Ptprc* (*Cd45*) labelling all immune cells and *Lyz2, Cd79a* and *Igkc* or *Cd3e* and *Trbc2*, to discriminate myeloid, B-cell and T-cell lineages, respectively. Cell type proportions were visualised using bar graphs in Prism (v9.3.1, GraphPad Software).

### Differential expression and gene ontology analysis

The FindMarkers function (using the Wilcoxon Rank Sum test) in Seurat was used to perform cell-specific differential expression analysis between wildtype and mutant cells, defining a differentially expressed gene as one with an average log_2_FC > 0.25 and adjusted *p*-value < 0.05. Results of differential expression were visualised using heatmaps or violin plots in Seurat and EnhancedVolcano (v1.12.0) plots. WT1 motif analysis was done using CiiiDER^18^, within 1kb base-pairs downstream and 500 base-pairs upstream of the transcriptional start site^6^, with stringency set to 0.1 and the WT1 motif (MA1627.1) defined by JASPAR 2022. Gene ontology (GO) analysis was performed by manually exporting lists of differentially expressed genes into the PANTHER webtool to group genes into GO biological processes^85^. Statistical overrepresentation (Fisher’s Exact test) was performed to calculate an enrichment score and false discovery rate (FDR) for each GO term, before plotting results in Prism.

### Ligand-receptor pair analysis

Inference of glomerular intercellular communication was performed using the NICHES package^37^. NICHES calculates the product of ligand expression by a given cell and the expression of its putative receptors in a pairwise manner, thus providing ligand-receptor analysis single-cell resolution. A NICHES object was created from a subset of the annotated scRNA-seq dataset containing only cells within the glomerular tuft, including podocytes and their physical or paracrine signalling candidates: glomerular endothelial cells, mesangial cells and parietal epithelial cells. The dataset was partitioned by genotype, before imputing ligand-receptor pairs for *Wt1*^*+/+*^ or *Wt1*^*R394W/+*^ glomerular cell types. A NICHES UMAP was created to visualise the pairwise interactions in two-dimensional space and the OmniPath database^86^ was used to create a list of putative ligand-receptor interactions between each cell pair. This list of putative ligand-receptor pairs was then manually curated, first by validating interactions on STRING^38^ (with “Co-occurrence” and “Text-mining” excluded to ensure interactions came from curated databases/experimental data only), followed by mining of the literature, to only include established intercellular interactions. The curated lists were used to generate heatmaps, listed in order of adjusted *p-*value and grouped by enrichment in *Wt1*^*+/+*^ glomeruli or *Wt1*^*R394W/+*^ glomeruli or those that were conserved across both.

### Cross-disease comparison of podocyte differential gene expression

To compare podocytes in *Wt1*^*R394W/+*^ glomerulopathy with other models of early glomerular disease, publicly accessible scRNA-seq data from Dynabead isolated glomeruli were used^9^.These included a nephrotoxic nephritis model (NTN, *n =* 2 mice and *n =* 2 controls) harvested at five days post injection in the absence of glomerulosclerosis. The second dataset was a leptin-deficiency (BTBR *Lep*^*ob/ob*^, *n =* 2 mutants and *n =* 2 controls) model of type 2 diabetes, taken at twelve weeks of age, characterised by proteinuria prior to glomerular lesions. The final dataset was a model of congenital nephrotic syndrome due to *Cd2ap* deficiency (*Cd2ap*^*−/−*^ *n =* 1 mouse or without *Cd2ap*^*+/+*^, *n =* 1 mouse) taken at three weeks, soon after the onset of proteinuria. Count matrices for each disease and their respective control groups were downloaded from the National Center for Biotechnology Information Gene Expression Omnibus (GSE146912). The quality control and pre-processing steps described above were applied separately to each dataset, to account for differences in data quality, batch effects and experimental conditions, such as murine genetic background. For each dataset, a further processing step of batch integration was performed using the Harmony package^87^ to ensure clustering of diseased and healthy podocytes within each condition. Podocytes were then identified by the co-expression of *Nphs1* and *Nphs2*. Differential expression analysis was run for healthy and diseased podocytes independently for each dataset, as above and the resulting lists were manually compared with that of *Wt1*^*R394W/+*^ glomerulopathy. The full script for the cross-disease comparison of scRNA-seq data has been made publicly available: https://github.com/daniyal-jafree1995/collaborations/blob/main/Chandleretal_2022_crossglomdiseasecomparison.R.

The number of differentially expressed genes present in each pathology were used to construct Venn diagrams. Genes, the upregulation or downregulation of which, was conserved by podocytes across all pathologies and those unique to *Wt1*^*R394W/+*^ glomerulopathy, were used to construct heatmaps. Each of the 23 conserved upregulated and 5 downregulated genes (found in all four analysed murine scRNA-seq datasets) then underwent targeted screening for similar dysregulation (fold-change > 0.25 and *p* < 0.05) in the Nephroseq database nephroseq.org of human microarray data, restricted to glomerular datasets only.

### Immunofluorescence

Immunofluorescence staining was performed on 7μm cryosections using primary antibodies against WT1 (1:100, ab89901, Abcam), F4/80 (MCA497G, Bio-Rad) and CD31 (1:400, MA3105, ThermoFisher) and imaged on a Zeiss Observer 7 Colour Fluorescence with Hamamatsu Flash 4.0v3 camera. The number of WT1^+^ cells found within the glomerular tuft was counted in 50 glomeruli per section and averaged for each individual. CD31^+^ staining was quantified as percentage coverage of tuft area in 50 glomeruli per section calculated in FIJI^88^ and averaged per animal. For myeloid counts, F4/80^+^ cells were counted within the glomerulus and in the field of vision outside the glomerulus at a consistent magnification, counts were taken over 50 glomeruli per animal and averaged for each mouse.

### Primary podocyte harvest and culture

For primary podocyte harvest, glomeruli were isolated from *Wt1*^*+/+*^ and *Wt1*^*R394W/+*^ mice at 4 weeks of age as previously described^89^. In brief kidneys were digested in DPBS^−/−^ with 1 mg/mL Collagenase A (C2674, Sigma) and 200 U/ml DNASe I (18047-019, Invitrogen) for 30 mins at 37°C with gentle agitation. Collagenase digested tissue was then sieved through a 100 μm cell strainer, collected by centrifugation and resuspended in 1 ml DPBS^−/−^. After washing, glomeruli were resuspended in growth media (RPMI-1640 (21875091, ThermoFisher) with 10% FBS, 1% Penicillin-Streptomycin and 1% Insulin-Transferrin-Selenium (41400045, ThermoFisher) and plated on Matrigel (354234, Corning) coated plates (diluted at 1:100). Glomeruli were incubated at 37°C, in 5% CO_2_ for seven days, at which point glomeruli were removed by 30 μm sieving and podocytes were left to fully differentiate for another 3 days. Podocyte total RNA was extracted (74136, Qiagen) and cDNA prepared using the gDNA Clear cDNA Synthesis Kit according to manufactures instructions (172-5035, Bio-Rad). Quantitative real-time polymerase chain reaction (RT-qPCR) was performed in duplicate using qPCRBIO SyGreen Mix Lo-ROX (PB20.11-05, PCRBioSystems) with gene specific primers for *Vegfa, Nrp1* and *Sulf1* standardised to *Gapdh*.

## Statistical methods

Statistical methods for scRNA-seq data have been described. All other statistical differences between *Wt1*^*+/+*^ and *Wt1*^*R394W/+*^ groups were analysed using Prism. Normality was assessed using the Shapiro-Wilk test and *t*-tests were used throughout. Statistical significance was accepted at *p* < 0.05 and graphed data is presented at mean ± standard deviation (SD).

## Data availability

Processed and raw data for scRNA-seq experiments will be uploaded to NCBI Gene Expression Omnibus database upon publication of the manuscript.

## Code availability

The codes generated during this study are available at Github repository here https://github.com/daniyal-jafree1995/collaborations.

## Acknowledgements

The authors would like to thank Dr Karin Straathof for her use of the 10X Chromium Controller and Dr Ayad Eddaoudi and the UCL Flow Cytometry Core Facility. This work was supported by a Wellcome Trust Investigator Award (220895/Z/20/Z, to D.A.L), a National Institute for Health Research (NIHR) Biomedical Research Centre at Great Ormond Street Hospital for Children NHS Foundation Trust and University College London Catalyst Fellowship (to J.C.), a Kidney Research Project Grant (Paed_RP_011_20170929, to A.M.W, D.A.L and J.C), a LifeArc/Great Ormond Street Children’s Charity research project grant (VS0322, to D.A.L, J.C and A.M.W), an MRC project grant (MR/T016809/1 to A.S.W.) and PhD studentships from Diabetes UK (17/0005733, to G.P, D.A.L) and the London Interdisciplinary Biosciences BBSRC funded Doctoral Training Partnerships (to S.M, D.A.L). D.J.J is supported by the UCL MB/PhD programme, a Child Health Research PhD studentship, a Rosetrees Trust PhD Plus award (PhD2020\100012) and a Foulkes Foundation Fellowship for postdoctoral research.

## Author contributions

J.C., D.A. L. and A.M.W. conceived and designed the project, J.C. conducted characterisation of the mouse line, optimised single-cell preparation, performed dynabead perfusions with M. KJ. and conducted the scRNA-seq protocol, with 10X machine support from A.P. J.C. isolated primary podocytes for transcript analysis and was responsible for podocyte culture and RT-qPCR analysis, J.C. conducted gene searching in Nephroseq. D.J., G.P. and A.M. performed all scRNA-seq bioinformatical analyses, in particular D.J. performed cell annotation, differential expression analyses, gene ontology and intercellular communication analyses and G.P. conducted WT1 motif analyses. J.C. manually curated cell-cell interaction outputs. S.M. and M.B. performed immunostaining, imaging and cell-counting analysis on murine tissue with support from W.M.. J.C. and D.J. collated and presented the figures and J.C., D.J. and D.A.L. wrote the manuscript with assistance from A.S.W and P.J.W..

## References

1. Ranganathan S. Pathology of Podocytopathies Causing Nephrotic Syndrome in Children. Front. Pediatr. 4, 32 (2016).

2. Cheong HI. Genetic tests in children with steroid-resistant nephrotic syndrome. Kidney Res. Clin. Pract. 39, 7–16 (2020).

3. Gnudi L, Benedetti S, Woolf AS, Long DA. Vascular growth factors play critical roles in kidney glomeruli. Clin. Sci. 129, 1225–1236 (2015).

4. Lennon R, Hosawi S. Glomerular cell crosstalk. Curr. Opin. Nephrol. Hypertens. 25 (2016).

5. Lefebvre J, et al. Alternatively spliced isoforms of WT1 control podocyte-specific gene expression. Kidney int. 88, 321–331 (2015).

6. Kann M, et al. Genome-Wide Analysis of Wilms’ Tumor 1-Controlled Gene Expression in Podocytes Reveals Key Regulatory Mechanisms. J. Am. Soc. Nephrol. 26, 2097–2104 (2015).

7. Hartwig S, et al. Genomic characterization of Wilms’ tumor suppressor 1 targets in nephron progenitor cells during kidney development. Development 137, 1189–1203 (2010).

8. He B, et al. Single-cell RNA sequencing reveals the mesangial identity and species diversity of glomerular cell transcriptomes. Nat. Commun. 12, 2141 (2021).

9. Chung J-J, et al. Single-Cell Transcriptome Profiling of the Kidney Glomerulus Identifies Key Cell Types and Reactions to Injury. J. Am. Soc. Nephrol. (2020).

10. Clark AR, et al. Single-Cell Transcriptomics Reveal Disrupted Kidney Filter Cell-Cell Interactions after Early and Selective Podocyte Injury. Am. J. Pathol. 192, 281–294 (2022).

11. Zambrano S, et al. Molecular insights into the early stage of glomerular injury in IgA nephropathy using single-cell RNA sequencing. Kidney int. 101, 752–765 (2022).

12. Gao F, et al. The Wt1+/R394W mouse displays glomerulosclerosis and early-onset renal failure characteristic of human Denys-Drash syndrome. Mol. Cell. Boil. 24, 9899–9910 (2004).

13. Little M, Wells C. A clinical overview of WT1 gene mutations. Hum. Mutat. 9, 209–225 (1997).

14. Korin B, Chung JJ, Avraham S, Shaw AS. Preparation of single-cell suspensions of mouse glomeruli for high-throughput analysis. Nat. Protoc. 16, 4068–4083 (2021).

15. Dumas SJ, et al. Phenotypic diversity and metabolic specialization of renal endothelial cells. Nat. Rev. Nephrol. 17, 441–464 (2021).

16. Zimmerman KA, et al. Single-Cell RNA Sequencing Identifies Candidate Renal Resident Macrophage Gene Expression Signatures across Species. J. Am. Soc. Nephrol. 30, 767–781 (2019).

17. Karaiskos N, et al. A Single-Cell Transcriptome Atlas of the Mouse Glomerulus. J. Am. Soc. Nephrol. 29, 2060–2068 (2018).

18. Gearing LJ, et al. CiiiDER: A tool for predicting and analysing transcription factor binding sites. PLoS One 14, e0215495 (2019).

19. Ishikawa T, et al. A novel podocyte protein, R3h domain containing-like, inhibits TGF-β-induced p38 MAPK and regulates the structure of podocytes and glomerular basement membrane. J. Mol. Med. 99, 859–876 (2021).

20. Fu J, et al. Regeneration of glomerular metabolism and function by podocyte pyruvate kinase M2 in diabetic nephropathy. JCI Insight 7 (2022).

21. Yuan Q, et al. Role of pyruvate kinase M2-mediated metabolic reprogramming during podocyte differentiation. Cell. Death. Dis. 11, 355 (2020).

22. Henique C, et al. Genetic and pharmacological inhibition of microRNA-92a maintains podocyte cell cycle quiescence and limits crescentic glomerulonephritis. Nat. Commun. 8, 1829 (2017).

23. Hiromura K, et al. Podocyte expression of the CDK-inhibitor p57 during development and disease. Kidney Int. 60, 2235–2246 (2001).

24. Barisoni L, Mokrzycki M, Sablay L, Nagata M, Yamase H, Mundel P. Podocyte cell cycle regulation and proliferation in collapsing glomerulopathies. Kidney Int. 58, 137–143 (2000).

25. Shankland SJ, Eitner F, Hudkins KL, Goodpaster T, D’Agati V, Alpers CE. Differential expression of cyclin-dependent kinase inhibitors in human glomerular disease: role in podocyte proliferation and maturation. Kidney Int. 58, 674–683 (2000).

26. Hosni ND, Anauate AC, Boim MA. Reference genes for mesangial cell and podocyte qPCR gene expression studies under high-glucose and renin-angiotensin-system blocker conditions. PLoS One 16, e0246227 (2021).

27. Chen Z, et al. A single-cell survey of the human glomerulonephritis. J. Cell. Mol. Med. 25, 4684–4695 (2021).

28. Foster MC, et al. Filtration markers as predictors of ESRD and mortality in Southwestern American Indians with type 2 diabetes. Am. J. Kidney. Dis. 66, 75–83 (2015).

29. Randles MJ, et al. Genetic Background is a Key Determinant of Glomerular Extracellular Matrix Composition and Organization. J. Am. Soc. Nephrol. 26, 3021–3034 (2015).

30. Roberts-Pilgrim AM, et al. Deficient degradation of homotrimeric type I collagen, α1(I)3 glomerulopathy in oim mice. Mol. Genet. Metab. 104, 373–382 (2011).

31. Connell LE, Helfman DM. Myosin light chain kinase plays a role in the regulation of epithelial cell survival. J. Cell. Sci. 119, 2269–2281 (2006).

32. Gorski M, et al. 1000 Genomes-based meta-analysis identifies 10 novel loci for kidney function. Sci. Rep. 7, 45040–45040 (2017).

33. McCarty G, Awad O, Loeb DM. WT1 protein directly regulates expression of vascular endothelial growth factor and is a mediator of tumor response to hypoxia. J. Biol. Chem. 286, 43634–43643 (2011).

34. Eremina V, et al. Glomerular-specific alterations of VEGF-A expression lead to distinct congenital and acquired renal diseases. J. Clin. Invest. 111, 707–716 (2003).

35. Wnuk M, Anderegg MA, Graber WA, Buergy R, Fuster DG, Djonov V. Neuropilin1 regulates glomerular function and basement membrane composition through pericytes in the mouse kidney. Kidney Int. 91, 868–879 (2017).

36. Herzog B, Pellet-Many C, Britton G, Hartzoulakis B, Zachary IC. VEGF binding to NRP1 is essential for VEGF stimulation of endothelial cell migration, complex formation between NRP1 and VEGFR2, and signaling via FAK Tyr407 phosphorylation. Mol. Biol. Cell. 22, 2766–2776 (2011).

37. Raredon MSB, Yang J, Kothapalli N, Kaminski N, Niklason LE, Kluger Y. Comprehensive visualization of cell-cell interactions in single-cell and spatial transcriptomics with NICHES. BioRxiv, 2022.2001.2023.477401 (2022).

38. Szklarczyk D, et al. The STRING database in 2021: customizable protein-protein networks, and functional characterization of user-uploaded gene/measurement sets. Nucleic Acids Res. 49, D605–d612 (2021).

39. Veron D, et al. Overexpression of VEGF-A in podocytes of adult mice causes glomerular disease. Kidney Int. 77, 989–999 (2010).

40. Sharma A, et al. A new role for the muscle repair protein dysferlin in endothelial cell adhesion and angiogenesis. Arterioscler. Thromb. Vasc. Biol. 30, 2196–2204 (2010).

41. Qi JH, et al. A novel function for tissue inhibitor of metalloproteinases-3 (TIMP3): inhibition of angiogenesis by blockage of VEGF binding to VEGF receptor-2. Nat. Med. 9, 407–415 (2003).

42. Koch S, et al. NRP1 presented in trans to the endothelium arrests VEGFR2 endocytosis, preventing angiogenic signaling and tumor initiation. Dev. Cell. 28, 633–646 (2014).

43. Vreeken D, Zhang H, van Zonneveld AJ, van Gils JM. Ephs and Ephrins in Adult Endothelial Biology. Int. J. Mol. Sci. 21 (2020).

44. Hellenthal KEM, Brabenec L NM. W. Regulation and Dysregulation of Endothelial Permeability during Systemic, Inflammation (2022).

45. Hino M, et al. Expression and regulation of adrenomedullin in renal glomerular podocytes. Biochem. Biophys. Res. Commun. 330, 178–185 (2005).

46. Dong N, Meng L, Xue R, Yu M, Zhao Z, Liu X. Adrenomedullin ameliorates podocyte injury induced by puromycin aminonucleoside in vitro and in vivo through modulation of Rho GTPases. Int. Urol. Nephrol. 49, 1489–1506 (2017).

47. Avraham S, Korin B, Chung JJ, Oxburgh L, Shaw AS. The Mesangial cell - the glomerular stromal cell. Nat. Rev. Nephrol. 17, 855–864 (2021).

48. Motegi S, Garfield S, Feng X, Sárdy M, Udey MC. Potentiation of platelet-derived growth factor receptor-β signaling mediated by integrin-associated MFG-E8. Arterioscler. Thromb. Vasc. Biol. 31, 2653–2664 (2011).

49. McGowan SE, McCoy DM. Neuropilin-1 and platelet-derived growth factor receptors cooperatively regulate intermediate filaments and mesenchymal cell migration during alveolar septation. Am. J. Physiol. Lung Cell. Mol. Physiol. 315, L102–l115 (2018).

50. Muhl L, et al. Neuropilin 1 binds PDGF-D and is a co-receptor in PDGF-D-PDGFRβ signaling. J. Cell. Sci. 130, 1365–1378 (2017).

51. Yeh CH, Chang CK, Cheng MF, Lin HJ, Cheng JT. The antioxidative effect of bone morphogenetic protein-7 against high glucose-induced oxidative stress in mesangial cells. Biochem. Biophys. Res. Commun. 382, 292–297 (2009).

52. Chan WL, Leung JC, Chan LY, Tam KY, Tang SC, Lai KN. BMP-7 protects mesangial cells from injury by polymeric IgA. Kidney Int. 74, 1026–1039 (2008).

53. Francavilla C, et al. The binding of NCAM to FGFR1 induces a specific cellular response mediated by receptor trafficking. J. Cell. Biol. 187, 1101–1116 (2009).

54. Gipson GR, Kattamuri C, Czepnik M, Thompson TB. Characterization of the different oligomeric states of the DAN family antagonists SOSTDC1 and SOST. Biochem. J. 477, 3167–3182 (2020).

55. Duan Z, et al. Specificity of TGF-β1 signal designated by LRRC33 and integrin α(V)β(8). Nat. Commun. 13, 4988 (2022).

56. Generous AR, et al. Trans-endocytosis elicited by nectins transfers cytoplasmic cargo, including infectious material, between cells. J. Cell. Sci. 132,(2019).

57. Li X, et al. Podocyte-specific deletion of miR-146a increases podocyte injury and diabetic kidney disease. Front. Med. 9, 897188 (2022).

58. Berthaud R, et al. Atypical severe early-onset nephrotic syndrome: Answers. Pediatr. Nephrol., (2022).

59. Natoli TA, et al. A mutant form of the Wilms’ tumor suppressor gene WT1 observed in Denys-Drash syndrome interferes with glomerular capillary development. J. Am. Soc. Nephrol.13, 2058–2067 (2002).

60. Shih NY, et al. Congenital nephrotic syndrome in mice lacking CD2-associated protein. Science. 286, 312–315 (1999).

61. Ju W, et al. Defining cell-type specificity at the transcriptional level in human disease. Genome Res. 23, 1862–1873 (2013).

62. Hodgin JB, et al. A molecular profile of focal segmental glomerulosclerosis from formalin-fixed, paraffin-embedded tissue. Am. J. Pathol. 177, 1674–1686 (2010)

63. Reich HN, et al. A molecular signature of proteinuria in glomerulonephritis. PLoS One 18;5(10):e13451 (2010)

64. Berthier CC, et al. Cross-species transcriptional network analysis defines shared inflammatory responses in murine and human lupus nephritis. J. Immunol. 189, 988–1001 (2012).

65. Peterson KS, et al. Characterization of heterogeneity in the molecular pathogenesis of lupus nephritis from transcriptional profiles of laser-captured glomeruli. J. Clin. Invest. 113, 1722–1733 (2004).

66. Woroniecka KI, Park AS, Mohtat D, Thomas DB, Pullman JM, Susztak K. Transcriptome analysis of human diabetic kidney disease. Diabetes 60, 2354–2369 (2011).

67. Ratelade J, et al. A murine model of Denys-Drash syndrome reveals novel transcriptional targets of WT1 in podocytes. Hum. Mol. Genet. 19, 1–15 (2010).

68. Schumacher VA, et al. WT1-dependent sulfatase expression maintains the normal glomerular filtration barrier. J. Am. Soc. Nephrol. 22, 1286–1296 (2011).

69. Takashima Y, et al. Heparan sulfate 6-O-endosulfatases, Sulf1 and Sulf2, regulate glomerular integrity by modulating growth factor signaling. Am. J. Physiol. Renal Physiol. 310, F395–408 (2016).

70. Ostalska-Nowicka D, Zachwieja J, Nowicki M, Kaczmarek E, Siwińska A, Witt M. Immunohistochemical detection of galectin-1 in renal biopsy specimens of children and its possible role in proteinuric glomerulopathies. Histopathology 51, 468–476 (2007).

71. Liu Y, et al. High glucose-induced Galectin-1 in human podocytes implicates the involvement of Galectin-1 in diabetic nephropathy. Cell. Biol. Int. 39, 217–223 (2015).

72. Brunskill EW, et al. Genes that confer the identity of the renin cell. J. Am. Soc. Nephrol. 22, 2213–2225 (2011).

73. Bennett MR, Czech KA, Arend LJ, Witte DP, Devarajan P, Potter SS. Laser capture microdissection-microarray analysis of focal segmental glomerulosclerosis glomeruli. Nephron Exp. Nephrol. 107, e30–40 (2007).

74. Potter AS, Drake K, Brunskill EW, Potter SS. A bigenic mouse model of FSGS reveals perturbed pathways in podocytes, mesangial cells and endothelial cells. PLoS One 14, e0216261 (2019).

75. Usui T, et al. Transcription factor MafB in podocytes protects against the development of focal segmental glomerulosclerosis. Kidney Int. 98, 391–403 (2020).

76. Chung AC, Lan HY. Chemokines in renal injury J. Am. Soc. Nephrol. 22, 802–809 (2011).

77. Michaelson JS, Wisniacki N, Burkly LC, Putterman C. Role of TWEAK in lupus nephritis: a bench-to-bedside review. J. Autoimmun. 39, 130–142 (2012).

78. Sasaki Y, Shimizu Y, Suzuki Y, Horikoshi S, Tomino Y. TWEAK/Fn14 system and crescent formation in IgA nephropathy. BMC Nephrol. 16, 27 (2015).

79. Sanchez-Niño MD, et al. Fn14 in podocytes and proteinuric kidney disease. Biochim. Biophys. Acta. 1832, 2232–2243 (2013).

80. Zheng GX, et al. Massively parallel digital transcriptional profiling of single cells. Nat. Commun. 8, 14049 (2017).

81. Kim D, Paggi JM, Park C, Bennett C, Salzberg SL. Graph-based genome alignment and genotyping with HISAT2 and HISAT-genotype. Nat. Biotechnol. 37, 907–915 (2019).

82. Hao Y, et al. Integrated analysis of multimodal single-cell data. Cell 184, 3573–3587.e3529 (2021).

83. McGinnis CS, Murrow LM, Gartner ZJ. DoubletFinder: Doublet Detection in Single-Cell RNA Sequencing Data Using Artificial Nearest Neighbors. Cell Systems 8, 329–337.e324 (2019).

84. Young MD, Behjati S. SoupX removes ambient RNA contamination from droplet-based single-cell RNA sequencing data. GigaScience 9, giaa151 (2020).

85. Mi H, et al. Protocol Update for large-scale genome and gene function analysis with the PANTHER classification system (v.14.0). Nat. Protoc. 14, 703–721 (2019).

86. Türei D, Korcsmáros T, Saez-Rodriguez J. OmniPath: guidelines and gateway for literature-curated signaling pathway resources. Nat. Methods 13, 966–967 (2016).

87. Korsunsky I, et al. Fast, sensitive and accurate integration of single-cell data with Harmony. Nat. Methods 16, 1289–1296 (2019).

88. Schindelin J, et al. Fiji: an open-source platform for biological-image analysis. Nat. Methods 9, 676–682 (2012).

89. Vasilopoulou E, et al. Loss of endogenous thymosin beta4 accelerates glomerular disease. Kidney Int. 90, 1056–1070 (2016).

