## Supplementary Figures for "Exploration of the single-cell transcriptomic landscape identifies aberrant glomerular cell crosstalk in a murine model of WT1 kidney disease"

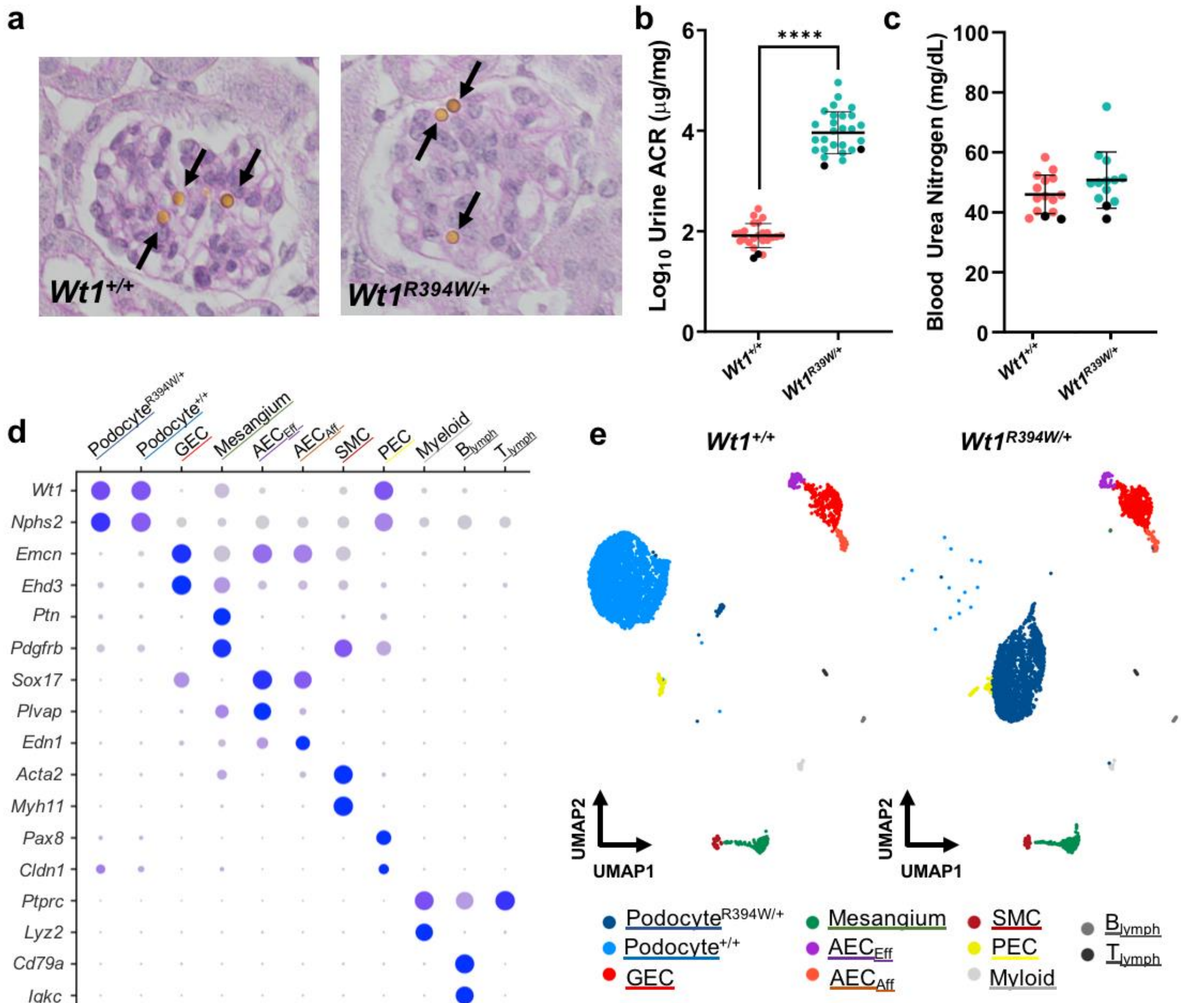


f

| Cell type | Mouse 1 *Wt1*$^{+/+}$ | Mouse 2 *Wt1*$^{+/+}$ | Mouse 3 *Wt1*$^{R394W/+}$ | Mouse 4 *Wt1*$^{R394W/+}$ |
|---|---|---|---|---|
| Podocyte$^{R394W/+}$ | 34 | 4 | 1280 | 757 |
| Podocyte$^{+/+}$ | 2365 | 760 | 10 | 5 |
| GEC | 202 | 96 | 297 | 126 |
| Mesangium | 79 | 41 | 118 | 79 |
| $AEC_{Eff}$ | 57 | 19 | 52 | 13 |
| $AEC_{Aff}$ | 50 | 23 | 42 | 14 |
| SMC | 38 | 10 | 39 | 12 |
| PEC | 27 | 14 | 27 | 10 |
| Myeloid | 25 | 10 | 23 | 3 |
| $B_{lymph}$ | 11 | 3 | 33 | 1 |
| $T_{lymph}$ | 14 | 8 | 9 | 6 |

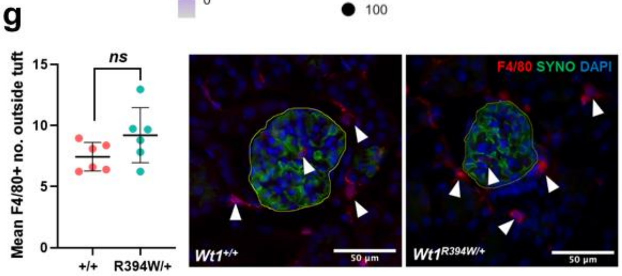


**Supplementary Figure 1: Supporting data for single-cell RNA seq analyses.** a) Glomeruli from *Wt1*$^{+/+}$ and *Wt1*$^{R394W/+}$ mice after Dynabead perfusion confirming bead penetration into *Wt1*$^{R394W/+}$ glomeruli at 4 weeks of age, prior to advanced glomerulosclerosis, confirming a non-biased harvest. b) $log_{10}$ albumin/creatinine ration (ACR) at 4 weeks of age with scRNA-seq donors highlighted in black, demonstrating their representative nature of the *Wt1*$^{R394W/+}$ mouse line. c) Blood urea nitrogen (BUN) levels at 4 weeks of age with scRNA-seq donors highlighted in black, demonstrating their representative nature. d) Dot blot of canonical marker genes used for cluster resolution for eleven cell type clusters. e) UMAPs split by *Wt1*$^{+/+}$ and *Wt1*$^{R394W/+}$ genotype, showing clear representation from both genotypes in each cell type cluster, with the exception of podocytes. f) Table of cell counts by cell type cluster and mouse donor, showing contribution from all four mice to each cluster, with the exception of podocytes. g) Extraglomerular myeloid (F4/80$^{+}$) cell counts, present in the area outside if the glomerular tuft in 4-week-old glomeruli (averaged over 50 glomeruli per animal), show no significant difference between *Wt1*$^{+/+}$ (7.48±0.48) and *Wt1*$^{R394W/+}$ (9.21±0.92) mice (*t*-test; $p = 0.1200$, $n = 6$ mice per group).

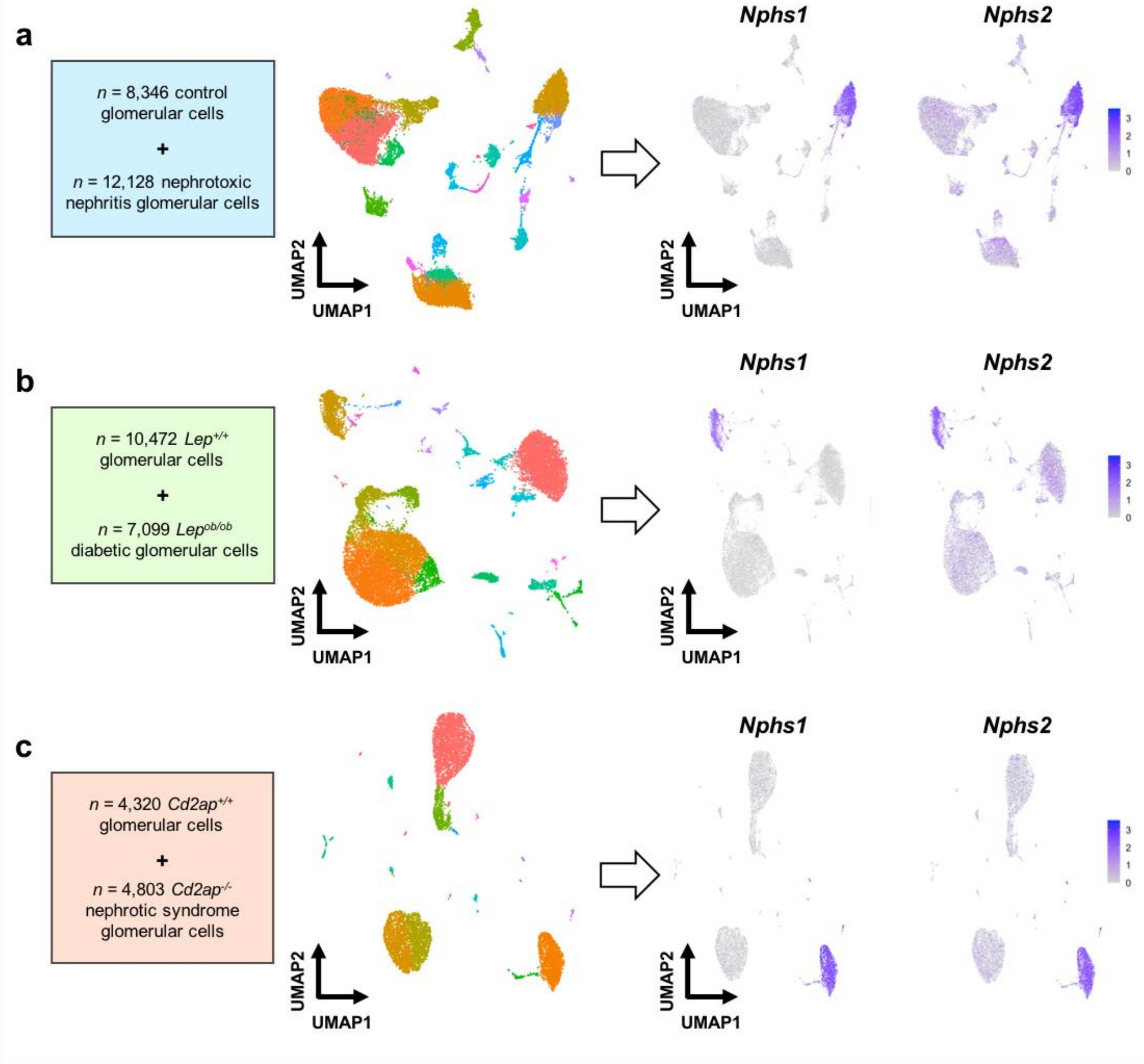


**Supplementary Figure 2: Isolation of podocytes from additional murine models of glomerular disease for cross-disease comparison**. Data were derived from the National Center for Biotechnology Information Gene Expression Omnibus (GSE146912)[9] and included: a) Two mice five days after induction with nephrotoxic nephritis (NTN, *n* = 1,450 podocytes) and two control mice (*n* = 1,491 podocytes). b) Two 12-week-old mice carrying a leptin mutation in homozygosity *(*$Lep^{ob/ob}$, *n* = 713 podocytes) modelling diabetic nephropathy and two littermate controls ($Lep^{+/+}$, *n* = 869 podocytes). c) One $Cd2ap^{-/-}$ mouse with congenital nephrotic syndrome (*n* = 423 podocytes) and one littermate control (*n* = 1337 podocytes). UMAPs were generated from each dataset independently and podocytes were identified by co-expression of $Nphs1^{+}$ and $Nphs2^{+}$. Differential expression analysis between healthy and diseased podocytes within each dataset served as an input for cross-disease comparison with the $Wt1^{R394W/+}$ model.

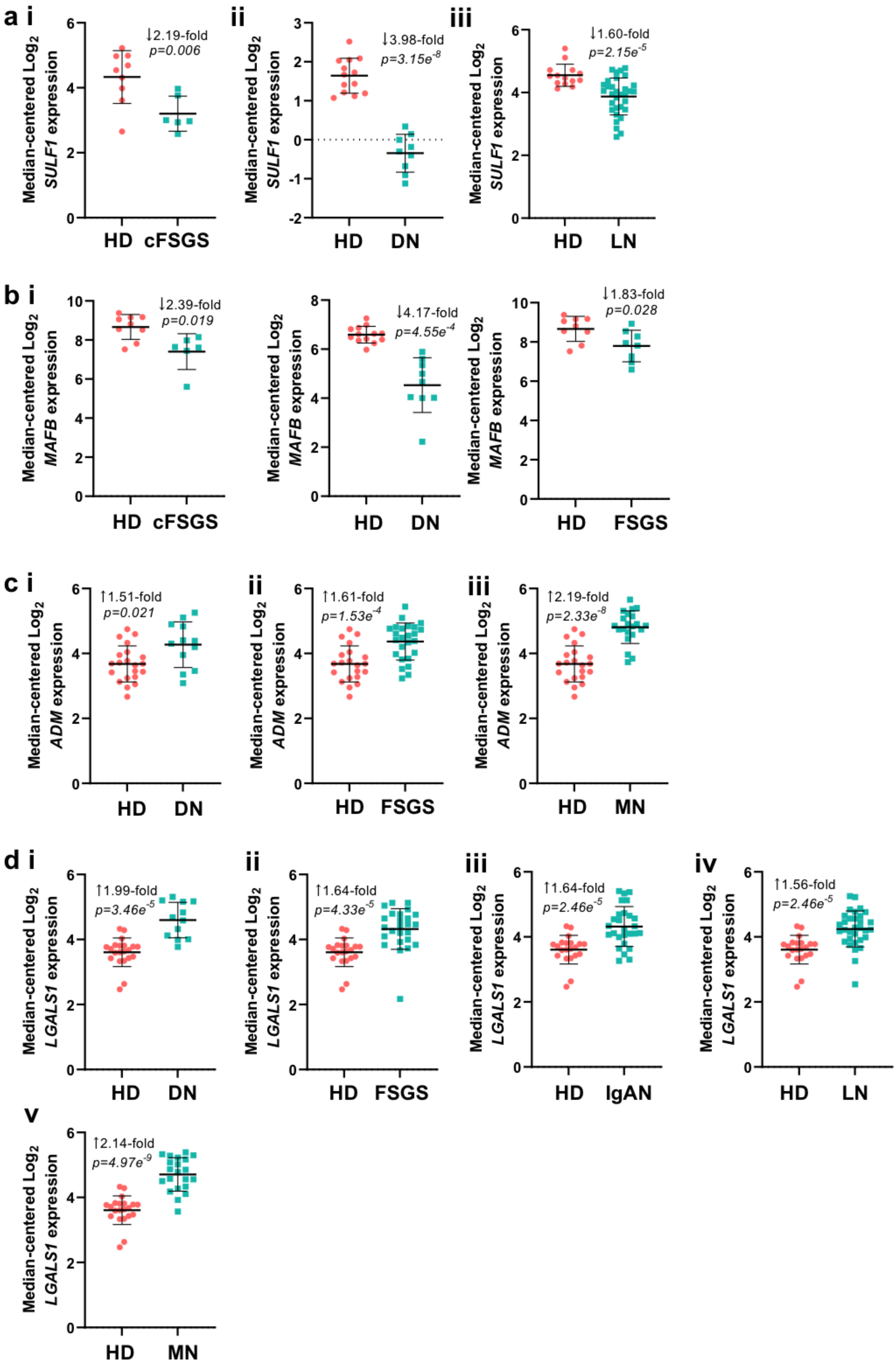
a i
Median-centered Log2 SULF1 expression
↓2.19-fold
p=0.006
HD
cFSGS
ii
↓3.98-fold
p=3.15e-8
HD
DN
iii
↓1.60-fold
p=2.15e-5
HD
LN
b i
Median-centered Log2 MAFB expression
↓2.39-fold
p=0.019
HD
cFSGS
↓4.17-fold
p=4.55e-4
HD
DN
↓1.83-fold
p=0.028
HD
FSGS
c i
Median-centered Log2 ADM expression
↑1.51-fold
p=0.021
HD
DN
ii
↑1.61-fold
p=1.53e-4
HD
FSGS
iii
↑2.19-fold
p=2.33e-8
HD
MN
d i
Median-centered Log2 LGALS1 expression
↑1.99-fold
p=3.46e-5
HD
DN
ii
↑1.64-fold
p=4.33e-5
HD
FSGS
iii
↑1.64-fold
p=2.46e-5
HD
IgAN
iv
↑1.56-fold
p=2.46e-5
HD
LN
v
↑2.14-fold
p=4.97e-9
HD
MN

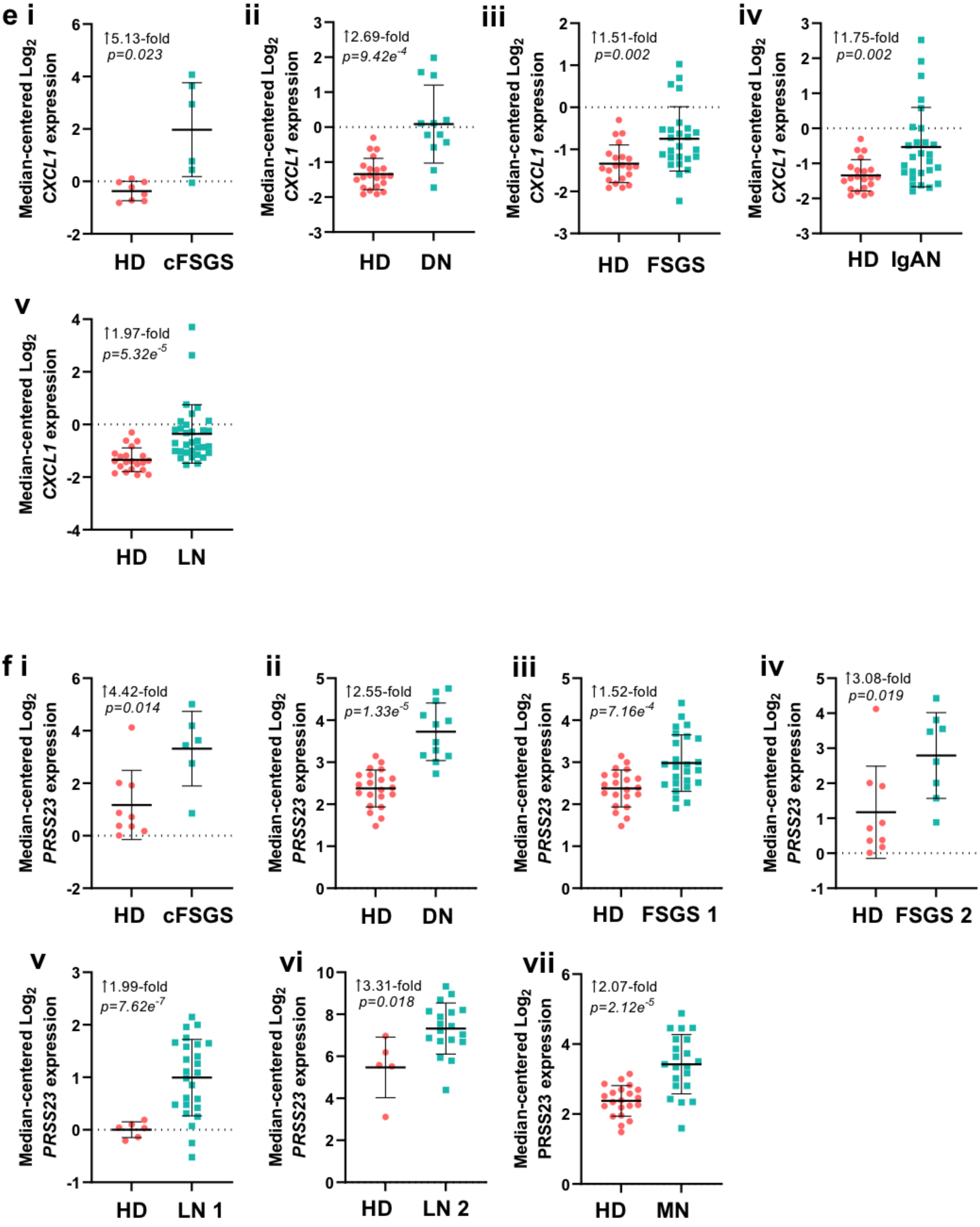
e i
↑5.13-fold
p=0.023
Median-centered Log2 CXCL1 expression
HD
cFSGS
ii
↑2.69-fold
p=9.42e-4
Median-centered Log2 CXCL1 expression
HD
DN
iii
↑1.51-fold
p=0.002
Median-centered Log2 CXCL1 expression
HD
FSGS
iv
↑1.75-fold
p=0.002
Median-centered Log2 CXCL1 expression
HD
IgAN
v
↑1.97-fold
p=5.32e-5
Median-centered Log2 CXCL1 expression
HD
LN
f i
↑4.42-fold
p=0.014
Median-centered Log2 PRSS23 expression
HD
cFSGS
ii
↑2.55-fold
p=1.33e-5
Median-centered Log2 PRSS23 expression
HD
DN
iii
↑1.52-fold
p=7.16e-4
Median-centered Log2 PRSS23 expression
HD
FSGS 1
iv
↑3.08-fold
p=0.019
Median-centered Log2 PRSS23 expression
HD
FSGS 2
v
↑1.99-fold
p=7.62e-7
Median-centered Log2 PRSS23 expression
HD
LN 1
vi
↑3.31-fold
p=0.018
Median-centered Log2 PRSS23 expression
HD
LN 2
vii
↑2.07-fold
p=2.12e-5
Median-centered Log2 PRSS23 expression
HD
MN

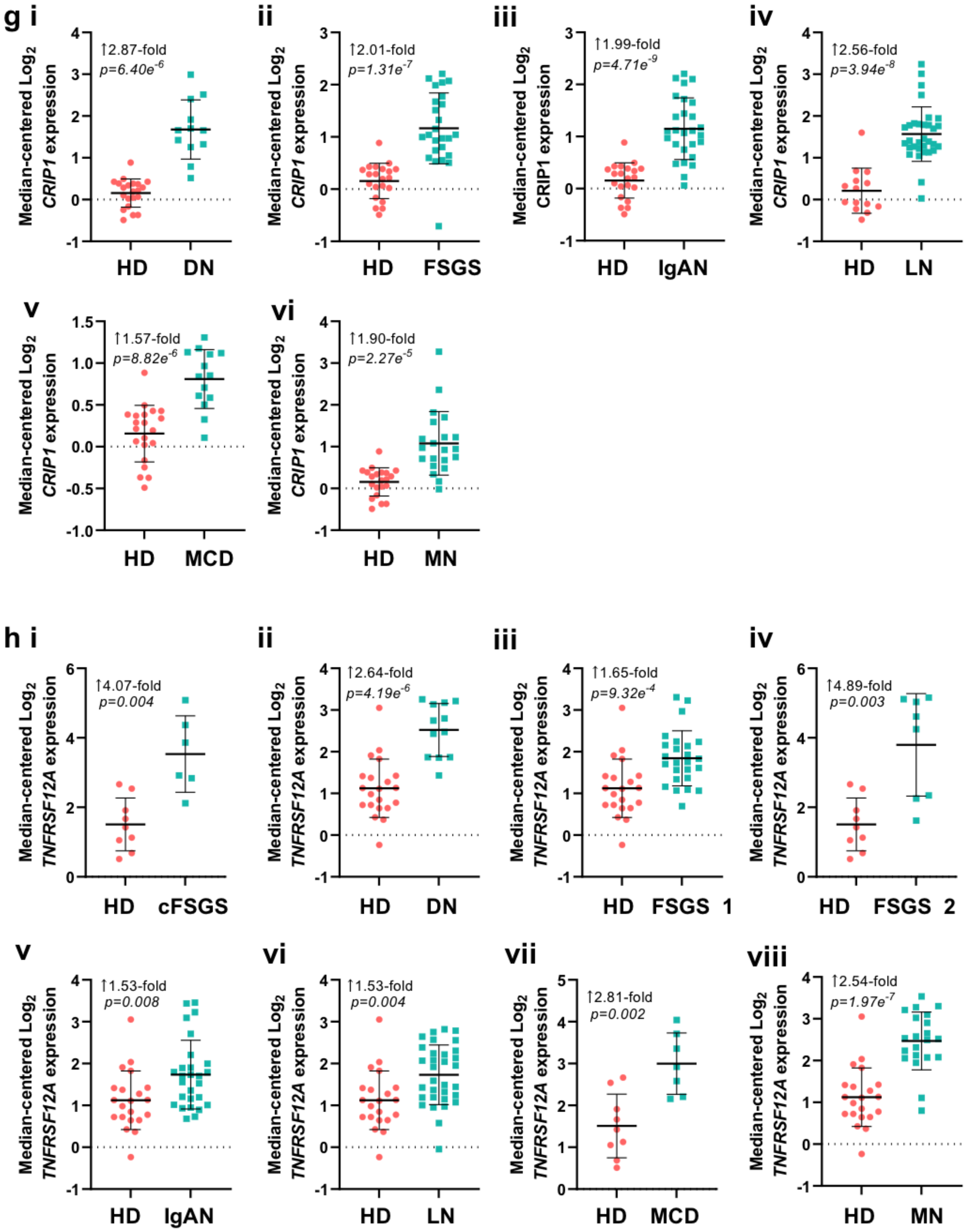


**Supplementary Figure 3: Dysregulated genes shown by individual sample for each human glomerular pathology (source Nephroseq)**. Fold-change and *p*-value indicated on each graph. Abbreviations as follows: HD, healthy donor; cFSGS, collapsing focal segmental glomerulosclerosis glomeruli; DN, diabetic nephropathy glomeruli; FSGS, focal segmental glomerulosclerosis glomeruli; IgAN, IgA nephropathy glomeruli; LN, lupus nephritis glomeruli; MCD, minimal change disease glomeruli; MN, membranous nephropathy glomeruli.

*See excel*

**Supplementary Table 1: Differentially expressed genes in glomerular cells in *Wt1*$^{R394W/+}$ mice**. Upregulated and downregulated genes in podocytes, glomerular endothelial cells, mesangial cells and parietal epithelial cells between *Wt1*$^{+/+}$ and *Wt1*$^{R394W/+}$ mice with an average $log_2FC > 0.25$ and adjusted $p$-value < 0.05. The presence of WT1 binding motifs, as predicted by CiiiDER[18], are indicated.

*See excel*

**Supplementary Table 2: Podocyte differentially expressed genes that are conserved across murine glomerular scRNA-seq datasets or unique to *Wt1*$^{R394W/+}$.** Podocyte genes conserved across three or more murine scRNA-seq datasets[9] of *Wt1*$^{R394W/+}$, nephrotoxic nephritis, *Lep*$^{ob/ob}$ diabetic nephropathy and *Cd2ap*$^{-/-}$, or only present in *Wt1*$^{R394W/+}$.

| Disease model | Up or Downregulated | No. of genes | Classified in GO:0002376 - *immune system process* |
|---|---|---|---|
| WT1 glomerulopathy (*Wt1*$^{R394W/+}$) | Upregulated | 3 unique (5) | ***Tmem178***, ***Prdx2,*** *Cxcl1*, *Cd59b*, ***Tcim*** |
| | Downregulated | 10 unique (14) | ***B2m***, *Zbtb20*, *Vegfa*, *Ctsh*, ***Tnfsf13***, ***Il34***, ***Bcl6***, ***Stat1***, ***H2-D1***, *H2-Q6*, ***H2-Q4***, ***H2-K1***, ***H2-T23***, ***Cd59a*** |
| Glomerulonephritis (nephrotoxic nephritis) | Upregulated | 12 unique (24) | *Cxcl2*, *Cd74*, *Cxcl13*, ***Ly96***, *Cxcl1*, ***Cxcl10***, ***Defb1***, ***Umod***, *Axl*, *Plvap*, ***Tmem176b***, ***Bst2***, ***H2-Eb1***, ***Mapkapk2***, *Lgals9*, ***Wfdc2***, ***H2-Ab1***, ***H2-Aa***, *Lgals3*, ***Ppp1r14b***, *Relb*, *Nfkb2*, ***Slamf9***, *Anxa1* |
| | Downregulated | 2 unique (7) | *Ifitm2*, *Zbtb20*, *Vegfa*, ***H2-Q7***, ***Prdx1***, *Ctsl*, *H2-Q6* |
| Diabetic nephropathy (BTBR *Lep*$^{ob/ob}$) | Upregulated | 18 unique (12) | ***Gm8909***, ***Akirin1***, ***Vegfd***, ***H2-Q10***, *Cxcl1*, ***Gata3***, *Axl*, *Plvap*, ***H2-DMb2***, *Lgals3*, ***Slpi***, ***Slc39a10*** |
| | Downregulated | 1 unique (7) | *Ifitm2*, *Zbtb20*, *Vegfa*, *Ctsh*, *Ctsl*, *H2-Q6*, ***Tir7*** |
| Congenital FSGS (*Cd2ap*$^{-/-}$) | Upregulated | 2 unique (11) | ***B2m***, *Cd74*, *Cxcl13*, *Cxcl1*, *Cd59b*, *Axl*, ***H2-D1***, *Lgals9*, *Relb*, |
| | Downregulated | 1 unique (1) | ***Ackr3*** |

**Supplementary Table 3: Podocyte differentially expressed genes classified in 'GO:0002376 - *immune system process'* across murine glomerular scRNA-seq datasets.** Podocyte differentially expressed genes from murine scRNA-seq datasets[9] of *Wt1*$^{R394W/+}$, nephrotoxic nephritis, *Lep*$^{ob/ob}$ diabetes and *Cd2ap*$^{-/-}$ that are classified into the broadest immunological GO term: *GO:0002376 - immune system process.* Genes unique to each disease model are in bold, the total number of differentially expressed genes classified is bracketed.
